# Spatio-temporally regulated maternal RNA decay factor essential for zebrafish development

**DOI:** 10.64898/2026.08.20.745964

**Authors:** Gopal Kushawah, Victor Daniel Aldas Bulos, Danielson Baia Amaral, Huzaifa Hassan, Stephanie H. Nowotarski, Ariel A. Bazzini

**Affiliations:** Stowers Institute for Medical Research, 1000 E 50th Street, Kansas City, MO 64110, USA; Department of Molecular and Integrative Physiology, University of Kansas Medical Center, 3901 Rainbow Blvd, Kansas City, KS 66160, USA

**Keywords:** Spatio-temporal maternal RNAs, RNA decay, CRISPR-Cas13d, CRISPR-Cas9, *Cth1*, Embryonic lethal, Gametogenesis, RNA-decay factor, Zebrafish, MZT

## Abstract

Maternal mRNA decay is essential for the **M**aternal-to-**Z**ygotic **T**ransition (MZT), yet its impact on spatial regulation has remained largely unexplored. We identify *Cth1* as the first maternally encoded RNA-decay factor regulated in a spatiotemporal manner outside the germline in zebrafish, driven by conserved 3′UTR *cis*-elements. RfxCas13d-mediated depletion of maternal *cth1* revealed early developmental defects that could not be assessed with SpyCas9 mutants due to infertility from impaired gametogenesis. Mechanistically, *Cth1* acts independently of zygotic genome activation, recognizing AU-rich motifs in maternal 3′UTRs to promote deadenylation and decay in a spatio-temporal manner; deletion of these motifs or depletion of *Cth1* stabilizes both reporter and endogenous transcripts. These findings establish *Cth1* as a key component of the maternal program and provide the first direct evidence that spatio-temporally regulated mRNA decay, outside the germline, contributes to early vertebrate development.

**Highlights:**

1. *Cth1* is a highly deposited maternal mRNA that undergoes rapid decay
2. 3’UTR of *Cth1* drives its spatio-temporal dynamics.
3. *Cth1* is essential for gametogenesis and early embryonic development.
4. CTH1 recognizes AU-rich elements in target 3′UTRs to trigger decay.

## Introduction

The maternal-to-zygotic transition (MZT) is a critical phase in eukaryotic embryogenesis characterized by the degradation of maternal messengers RNAs (mRNAs) and transcriptional activation of the zygotic genome (Giraldez et al, 2006; Kojima et al, 2025; Vastenhouw et al, 2019; Walser & Lipshitz, 2011). Initially, maternal mRNAs deposited by the female into the oocyte prior to fertilization regulate early embryonic development. Timely clearance of these maternal mRNAs is essential to ensure proper embryogenesis. Mechanisms regulating the decay of maternal mRNAs can be classified based on whether they are dependent on zygotic transcription. A maternally directed program functions independently or prior to zygotic transcription, whereas a zygotically directed program requires active zygotic transcription (Brantley & Di Talia, 2024; Giraldez et al, 2006; Kojima et al, 2025; Kontur et al, 2020). Thus, maternal RNA clearance pathways may originate either from maternal cytoplasmic factors inherited within the oocyte or from factors synthesized *de novo* from the zygotic genome (Kojima et al, 2025).

In different vertebrates, the zygotic program includes microRNAs, such as miR-430 in zebrafish or miR-427 in *Xenopus*, which are zygotically expressed and facilitate the clearance of many maternal and zygotic mRNAs (Baia Amaral et al, 2024; Bazzini et al, 2012; Giraldez et al, 2006; Lund et al, 2009). In contrast, in zebrafish and *Xenopus* embryos the maternal program utilizes codon optimality, where translation, influenced by codon usage, directly impacts mRNA stability (Bazzini et al, 2016; Mishima & Tomari, 2016). Additional maternal clearance pathways involve RNA-binding proteins that recognize distinct *cis*-elements. Examples include YTHDF (M^6^A modification) and TUT4/7 (Uridylation) in zebrafish (Chang et al, 2018; Kontur et al, 2020; Zhao et al, 2017), Smaug and BRAT in *Drosophila* (Laver et al, 2015; Tadros et al, 2007), EDEN-BP/Celf1 in *Xenopus* (Paillard et al, 1998), BTG4 and PABPN1L in mouse (Liu et al, 2016; Yu et al, 2016; Zhao et al, 2020). The ZFP36L family of proteins, which target AU-rich elements in 3′UTRs, have been implicated in mRNA clearance during mouse oocyte maturation leading to female infertility (Ball et al, 2014; Ramos, 2012). However, their contribution to mRNA clearance during the MZT remains unclear (Sha et al, 2018). AU-rich elements associated with unstable mRNAs have been reported across species, including *Xenopus* (Audic et al, 1998), *Drosophila* (Choi et al, 2014) and zebrafish (te Kronnie et al, 1999; Vejnar et al, 2019). However, the identity and functional relevance of the factors recognizing these elements during early embryogenesis remain largely unresolved.

Most studies on maternal mRNA degradation in zebrafish have relied on bulk mRNA sequencing, treating the embryo as a homogeneous entity and focusing on temporal dynamics (Baia Amaral et al, 2024; Bazzini et al, 2016; Giraldez et al, 2006; Vejnar et al, 2019). While these approaches have deepened our understanding of the timing of maternal mRNA clearance, they do not provide insight into potential spatial differences in mRNA distribution. Emerging single-cell and spatial transcriptomic methods offer new opportunities to study RNA accumulation with spatial resolution. However, the functional significance of spatially distinct mRNAs and the mechanisms underlying their spatial distribution during early development remain largely unexplored (Fishman et al, 2024; Holler et al, 2021; Satija et al, 2015; Wan et al, 2026).

In many cases where mRNAs show spatially restricted patterns, localized zygotic transcription can often explain the cell specific accumulation (Das et al, 2021; Martin & Ephrussi, 2009). However, for maternal mRNAs, differential spatial degradation provides an alternative mechanism for generating regional mRNA accumulation. Indeed, earlier studies in zebrafish, *Drosophila*, and sea urchin have demonstrated spatially regulated maternal mRNA decay resulting in maternal mRNAs accumulation in the germline (Gavis & Lehmann, 1992; Koprunner et al, 2001; Oulhen et al, 2013; Pamula & Lehmann, 2024). In zebrafish, for example, *nanos1* and *tdrd7* mRNAs prevent miR-430-mediated degradation in primordial germ cells due to protection by the germline-specific RNA-binding protein Dead end 1 (*Dnd1*), resulting in localized mRNA stability (Kedde et al, 2007; Mishima et al, 2006). However, the broader functional significance of spatially regulated maternal mRNA localization and particularly for genes outside of germline contexts, remains largely unexplored. This gap in knowledge is due in part to the historical lack of tools capable of efficiently and selectively manipulating maternal RNAs in early vertebrate embryos.

Here, we exploit new approaches to identify *cth1* as a previously uncharacterized, maternally deposited mRNA decay factor in zebrafish. Phylogenetic analyses and previous studies indicate that *cth1* has conserved orthologs across yeast, fish, and mammals, including the vertebrate ZFP36L gene family (Makita et al, 2021; Martinez-Pastor et al, 2013; Stevens et al, 1998; Treguer et al, 2013). These orthologs have been shown to promote mRNA decay by binding AU-rich elements in 3′UTRs in yeast, *Xenopus*, mouse and humans, implicating them in diverse RNA metabolic pathways (Baou et al, 2011; Barlit et al, 2024; Bye et al, 2018; Cicchetto et al, 2023; Cook et al, 2022; Rynne et al, 2023). In zebrafish, we found that *cth1* exhibits spatiotemporal instability driven by its 3′UTR. Using CRISPR-RfxCas13d and SpyCas9, we reveal its essential roles in maternal mRNA clearance during the MZT and in gametogenesis. Further, *Cth1* mediates the deadenylation and decay of mRNAs through AU-rich elements in their 3′UTRs and functions independently of zygotic transcription. This study establishes *Cth1* as a maternal factor regulating mRNA stability prior and independent of the zygotic genome activation, and as the first mRNA shown to display spatiotemporal dynamics due to stability mechanisms outside the germline.

## Results

### *Cth1* is one of the most highly maternally deposited mRNAs but is rapidly degraded during the maternal-to-zygotic transition

Maternal mRNA decay has predominantly been studied in a temporal dimension, with the strong evidence for spatial regulation in vertebrate embryos coming from germ cells, where selected maternal transcripts are protected while the same transcripts decay in neighboring somatic cells (Koprunner et al, 2001; Mishima et al, 2006; Oulhen et al, 2013). Whether spatially restricted maternal mRNA decay/protection also occurs beyond germ cells, and whether it contributes to early embryonic development, remains unsolved. To search for maternal RNAs regulated in this manner outside the germline, we analyzed single-cell/spatial transcriptomics and SLAM-seq (Baia Amaral et al, 2024; Satija et al, 2015). From the initial analysis of these datasets, we narrowed down to *Cth1*. Ribo-Zero and poly(A)-enriched RNA-seq analyses across early embryogenesis identified *cth1* as one of the most abundant maternal transcripts, ranking within the top 75 in both ribosomal RNA–depleted and poly(A)-enriched datasets (Figs. 1A and S1A) [45]. After fertilization, a substantial fraction of maternal mRNAs exhibits short poly(A) tails (Chang et al, 2018), as indicated by their lower abundance in poly(A)-enriched libraries compared to total

**Figure 1.**
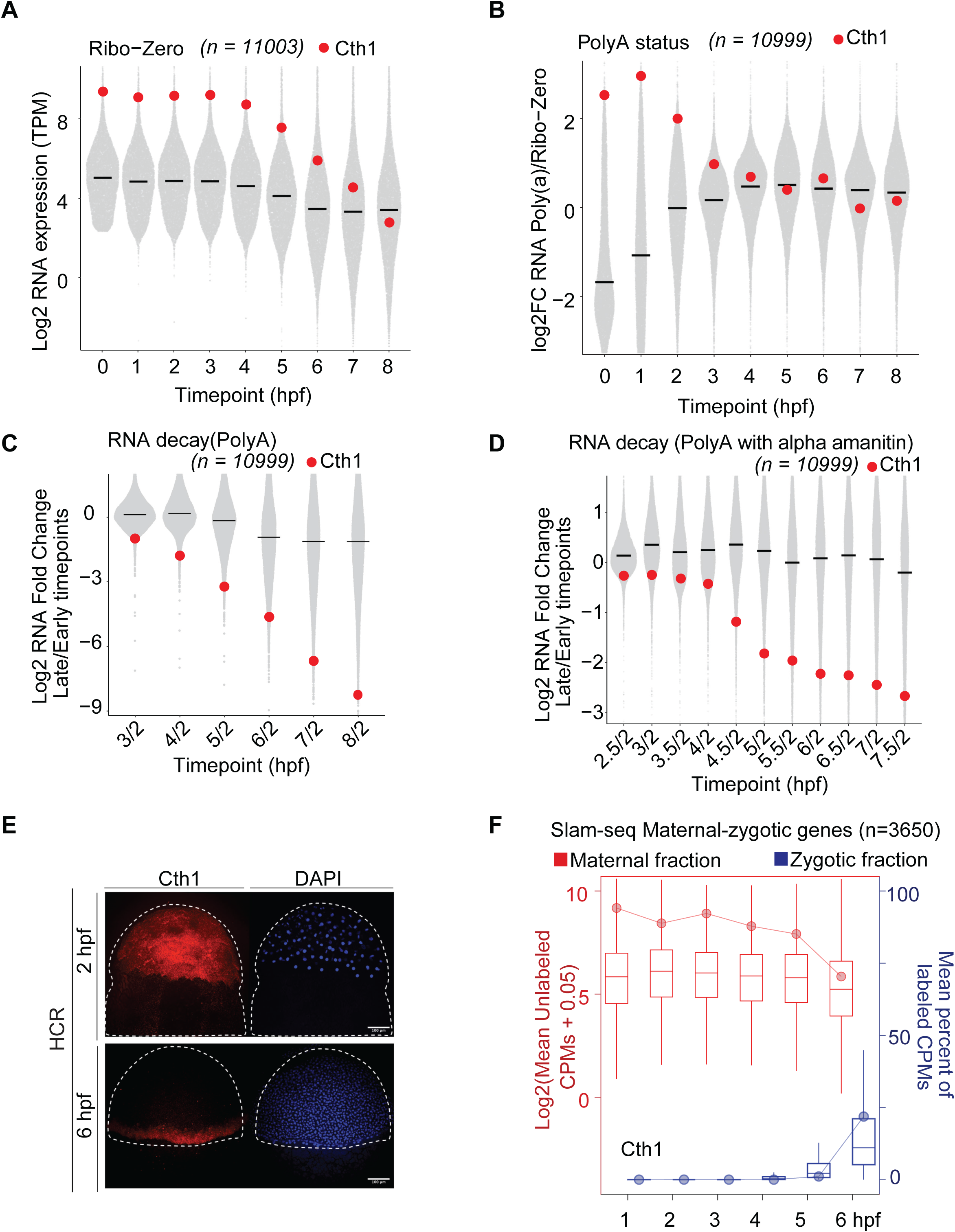
*Cth1* is one of the most highly maternally deposited mRNAs but is rapidly degraded during the maternal-to-zygotic transition. A. Ribosomal RNA depleted RNA sequencing (Ribo-zero) time course (0-8 hpf) showing *cth1* mRNA is among the top maternally deposited (top 75) mRNA at 0 hpf while getting rapidly degraded with time by 8 hpf. Data is derived from SG Medina-Muñoz et al., *Genome Biology,* 2021. B. Poly(A)-status measured as the fold change between PolyA and Ribo-zero RNA sequencing at different time points showing that *cth1* is one of the maternal RNA provided with already extended Poly(A)-tail (0 hpf) while getting deadenylated over time. Data is derived from SG Medina-Muñoz et al., *Genome Biology,* 2021. C. mRNA decay measured as fold change between PolyA RNA sequencing at 2 hpf and indicated time point (2-8 hpf) showing that *cth1* mRNA is one of the most unstable mRNA (top 20). Data is derived from SG Medina-Muñoz et al., *Genome Biology,* 2021. D. mRNA decay in the absence of zygotic expression measured as fold change between polyA RNA sequencing at 2 hpf and indicated time point (2.5-7.5 hpf) from embryos injected with α-amanitin showing that *cth1* mRNA degradation is maternally programmed independent of zygotic transcription. Data is derived from SG Medina-Muñoz et al., *Genome Biology,* 2021. E. Whole mount Hybridization Chain Reaction (HCR) showing spatio-temporal *cth1* mRNA (red) distribution and DAPI (blue) staining at 2 and 6 hpf zebrafish embryos. Scale bar, 100 μm. F. SLAM-seq time course (1-6 hpf) showing the maternal (unlabeled) and zygotic (labeled) contribution for *cth1* indicating that even at 6 hpf the maternal contribution is predominant compared to the zygotic. Data is derived from DB Amaral et al., *Genome Biology,* 2024.

RNA, also known as poly(A)-tail status (Fig. 1B) (Medina-Munoz et al, 2021). These mRNAs typically undergo polyadenylation within the first ∼2 hours post-fertilization (hpf) (Fig. 1B), as clearly illustrated by comparing poly(A)-selected mRNA levels against total RNA depleted of ribosomal RNA (Medina-Munoz et al, 2021). Interestingly, *cth1* mRNAs are maternally provided with extended poly(A)-tails, exhibiting one of the highest poly(A)-tail status at 0 hpf (top 75) (Fig. 1B).

Despite its high initial abundance and high poly(A)-tail status, *cth1* mRNA is remarkably unstable (top 20), rapidly decaying within the first 8 hpf (Fig. 1C, Fig. S1A). This rapid degradation is evidenced by pronounced fold changes relative to 0 hpf in Ribo-Zero (Fig. S1B) and to 2 hpf in poly(A)-selected RNA-seq datasets (Fig. 1C). To further characterize *cth1* dynamics, we analyzed SLAM-seq data during early embryogenesis (Baia Amaral et al, 2024). This technique distinguishes newly synthesized mRNAs from pre-existing ones by incorporating a uridine analog into nascent RNAs, which are later identified as T>C mutations during sequencing (Baia Amaral et al, 2024; Bhat et al, 2023; Herzog et al, 2017). SLAM-seq allows gene-specific quantification of both labeled (zygotic) and unlabeled (maternal) mRNAs and provides direct measurements of mRNA stability based on the persistence of unlabeled RNA. Using this approach, we confirmed that *cth1* is highly unstable, as reflected by its rapid loss from the unlabeled fraction (Fig. S1C).

The instability of *cth1* persisted in embryos injected with α-amanitin, indicating that its degradation is maternally programmed and independent of zygotic factors (Fig. 1D). Consistent with this observation, *cth1* degradation was also unaffected in Maternal-Zygotic-dicer mutants (lacking functional miR-430) demonstrating that its decay is independent of the miR-430 pathway (Fig. S1D). Ribosome profiling further reveals high ribosome occupancy and translational efficiency for *cth1* during early development (Fig. S1E), a pattern characteristic of transcripts previously shown to play critical roles in early embryogenesis (da Silva Pescador et al, 2024; Lee et al, 2013) and suggesting an important early function for Cth1.

Interestingly, despite rapid global degradation, *cth1* mRNA exhibited distinct localization to the embryo margin at 6 hpf based on whole-mount Hybridization Chain Reaction (HCR) assays (Fig. 1E) and whole embryo reconstruction based on single cells sequence and weMERFISH (Satija et al, 2015; Wan et al, 2026) (Fig. S1F). HCR assays revealed a uniform distribution at 2 hpf, followed by clear marginal enrichment at 6 hpf (Fig. 1E). These results raised two potential hypotheses: (i) maternal *cth1* mRNAs are entirely degraded, and cell-specific zygotic transcription drives marginal localization, or (ii) maternal *cth1* undergoes spatially regulated degradation (or cell-specific stabilization), resulting in marginal enrichment. Supporting the latter hypothesis, SLAM-seq analysis revealed a predominance of unlabeled *cth1* mRNAs at 6 hpf, indicating maternal origin rather than new transcription (Fig. 1F). Collectively, these data strongly suggest that the marginal localization of *cth1* mRNA results from spatially regulated maternal mRNA decay rather than zygotic transcription.

### The *cth1* 3′UTR harbors *cis*-regulatory elements driving marginal localization

Inspired by the ability of the nanos 3′UTR to mediate germline localization during embryogenesis (Koprunner et al, 2001), we tested whether the *cth1* marginal localization depends on *cis*-regulatory elements within its 3′UTR. We co-injected single-cell stage zebrafish embryos with TagRFP mRNA as an internal control and GFP mRNA fused either to the *cth1* 3′UTR or the control 3′UTR (non-zebrafish 3′UTR sequence) (Fig. 2A). Using HCR, we visualized the ectopic mRNA distribution at 2.5, 4, 5, and 6 hpf (Fig. 2B). GFP mRNA containing the *cth1* 3′UTR exhibited robust marginal localization at 5 and 6 hpf, whereas TagRFP and GFP fused to the control 3′UTR remained uniformly distributed (Fig. 2B). These findings suggest the presence of *cis*-regulatory elements within the *cth1* 3′UTR sufficient to drive marginal enrichment. The encoded GFP protein (signal) also accumulated in the same marginal domain, suggesting that the *cth1* 3′UTR directs both transcript localization and local protein production during gastrulation (Fig. 2C).

**Figure 2.**
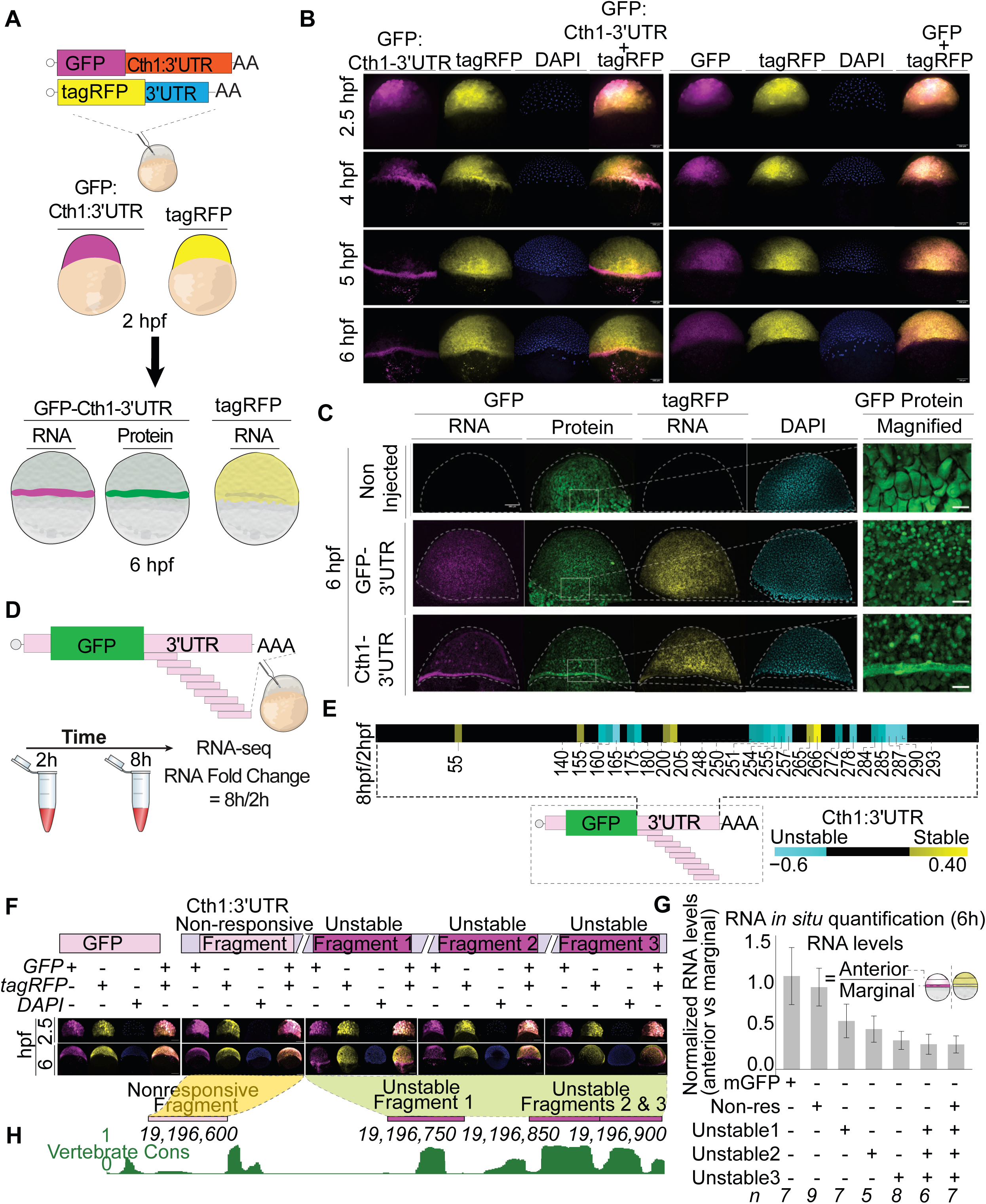
The *cth1* 3′UTR harbors *cis*-regulatory elements driving marginal localization. A. Schematic illustration of the injection strategy used to assess mRNA localization and protein distribution at 6 hpf. mRNAs encoding GFP containing either the *cth1* 3′UTR or a control 3′UTR were co-injected with a tagRFP mRNA as an injection control. B. GFP and TagRFP HCR showing that all injected mRNAs were uniformly distributed at 2 hpi, but then, only GFP-*cth1*-3’UTR mRNA was localized to the marginal region over time, while the GFP-*β-globin* 3′UTR and TagRFP mRNA remain uniformly distributed across the embryo. Scale bar, 100 μm. C. HCR combined with fluorescence imaging showing mRNA (magenta) and protein (green) localization at 6 hpf. GFP-*cth1*-3′UTR mRNA and its encoded protein are preferentially enriched at the embryonic margin, whereas GFP fused to a control 3′UTR (non-zebrafish sequence) shows uniform distribution throughout the embryo. The co-injected tagRFP mRNA (yellow) remains uniformly distributed. The rightmost panel shows a magnified view of GFP protein localization at the margin. Non-injected embryos were included as controls; residual green signal reflects autofluorescence from the yolk, which is detectable due to the high exposure times which is required to image low GFP levels at this stage. DAPI (cyan) marks nuclei. Scale bar, 100 μm. D. Schematics represents the tilling array of 50 nt *cth1* 3’UTR fragments library injections, the stability of the of fragments was assayed by comparing library-specific RNA-seq from embryos collected at 2 and 8 hpi. E. Heat map showing the stability of different *cth1* 3’UTR fragments between 8 and 2 hpi. Cyan color indicates unstable regions, black color non-responding region and yellow stable regions. The number represent the 5’position of the fragment with respect to *cth1* 3’UTR. F. GFP-*control* 3′UTR and TagRFP HCR showing that all injected mRNAs were uniformly distributed at 2.5 hpi, but then, only GFP-containing the unstable *cth1*-3’UTR 50 nt fragments (unstable, 1, 2, and 3) identified by the tilling library gets enriched to the marginal region at 6 hpi, while the GFP containing one stable *cth1*-3’UTR 50 nt fragment (tilling array library) and other injected control TagRFP mRNA display uniform distribution across the embryo. Scale bar, 200 μm. G. Bar plot showing normalized HCR RNA quantification for GFP vs TagRFP RNAs between two regions of the same embryos (anterior vs. marginal) for different *cth1* 3′UTR fragments and controls. *n* represents the number of embryos analyzed. H. Conservation across vertebrates UCSC tracks showing that the three unstable fragments of c*th1* 3’UTR are conserved (green color).

To further dissect these *cis*-regulatory regions within the *cth1* 3′UTR, we generated a tiling library consisting of 80 overlapping 50-nucleotide (nt) fragments from the *cth1* 3′UTR (≤5-nt sliding windows) (Fig. 2D). Each fragment was cloned downstream of GFP to mimic the full-length construct. As a control, we included 20 fragments spanning the *sod1* 3′UTR, known to contain a functional miR-430 target site. Following injection of the mRNA library into single-cell embryos, RNA was collected at 2- and 8-hours post-injection for specific library sequencing (Fig. 2D). As expected, *sod1* fragments containing *miR*-430 target sites displayed pronounced instability over time, validating our approach for identifying destabilizing elements (Figs. S2A, S2B). In the *cth1* 3′UTR, we identified three regions where fragments showed significant downregulation over time (Figs. 2E, S2C).

We next examined whether these destabilizing fragments were sufficient to drive marginal localization. Three 50-nt fragments containing each *cth1* 3′UTR destabilizing fragment and a *cth1* 50-nt non-responsive fragment were individually cloned as 3′UTR of GFP mRNA and injected into single-cell embryos alongside TagRFP as an internal control (Fig. 2F). At 2 hpf, all injected mRNAs were uniformly distributed (Fig. 2F). By 6 hpf, however, GFP mRNAs containing the individual destabilizing fragments displayed different degrees of RNA decay leading to marginal patterning (Figs. 2F-G and S2D), whereas non-responsive fragments and control mRNAs (TagRFP) remained uniformly distributed (Figs. 2F-G, S2D). Interestingly, these destabilizing regions were conserved across fish species (Fig. 2H). Consistent with the individual fragment results, combining all three destabilizing regions into a single 175-nt fragment produced an additive effect on decay and marginal localization (Figs. 2G, S2D-E).

Together, our results indicate that these three conserved 3′UTR regions contain *cis*-regulatory elements that contribute to regulation of spatiotemporal mRNA decay, thereby driving marginal localization of *cth1*. The evolutionary conservation underscores the potential significance of this regulatory mechanism across fish species.

### Cth1 F0 mutants do not exhibit early developmental defects but display severe infertility

Given that *cth1* localization during early embryogenesis occurs independently of its zygotic expression (Fig. 2) and that the zygotic contribution of *cth1* is minimal at early developmental stages (Fig. 1F), we hypothesized that F0 *cth1* mutants might lack early developmental phenotypes. To test this, we generated F0 *cth1* mutants by co-injecting SpyCas9 mRNA with either individual or combined guide RNAs (gRNAs) targeting the *cth1* coding region (Fig. 3A). We confirmed high mutagenesis efficiency (>85%) at 24 hpf by sequencing all targeted *cth1* loci along with the *albino* gene, which was also targeted independently as a positive control (Fig. 3B). Embryos injected with SpyCas9 and gRNAs targeting albino displayed the expected pigmentation loss (Fig. S3A). Consistent with our hypothesis, despite the high mutagenesis rate (Fig. 3A), no developmental defects were observed in *cth1* F0 mutant embryos at 6 and 24 hpf compared to SpyCas9-alone or uninjected controls (SpyCas9 alone, gRNA and albino) (p ≥ 0.15, SpyCas9 alone vs SpyCas9 plus each gRNA, chi-square test) (Fig. 3C, S3B). This suggests that early embryonic development does not significantly rely on the low zygotic *cth1* expression.

**Figure 3.**
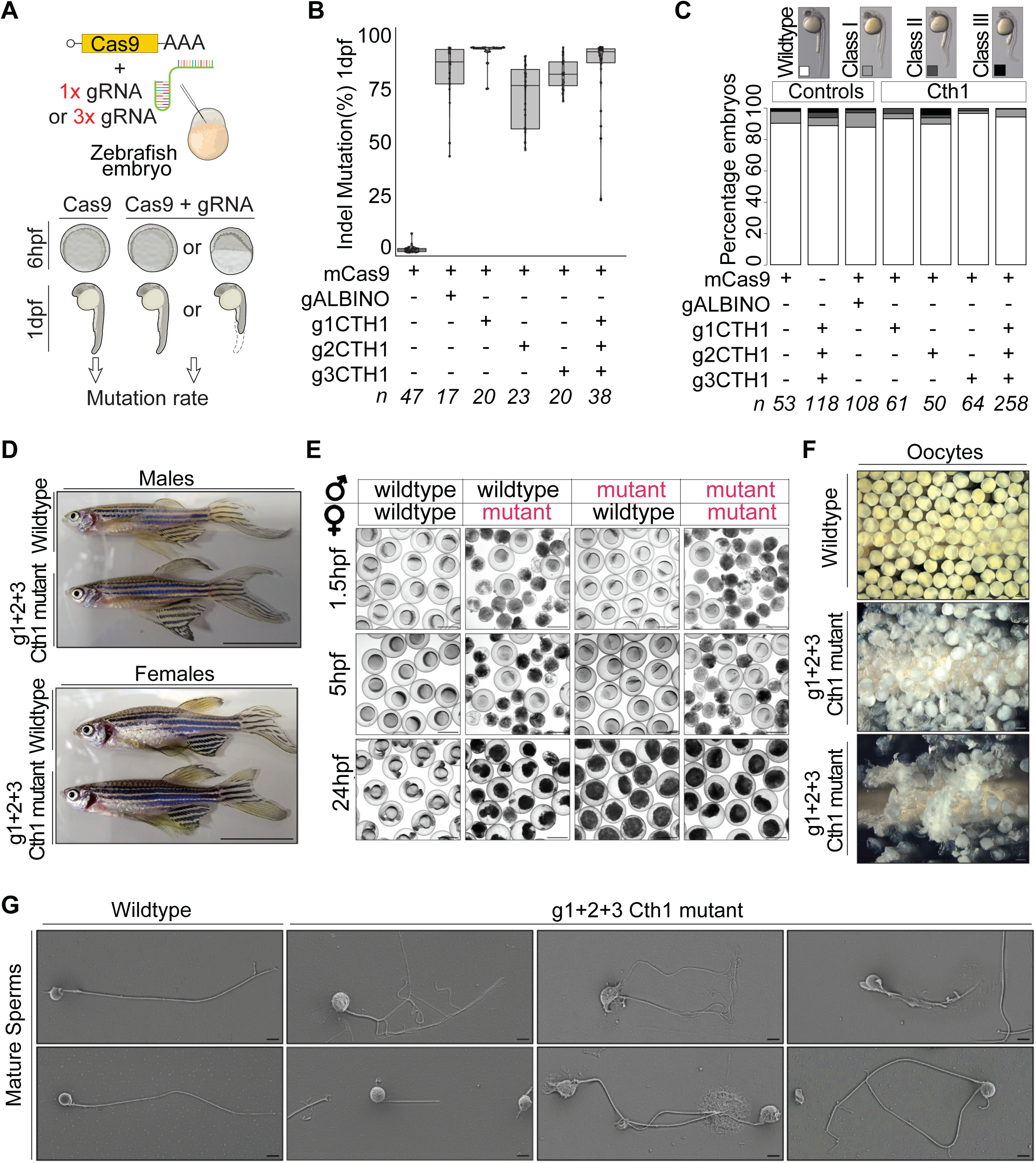
*Cth1* F0 mutants do not exhibit early developmental defects but display severe infertility. A. Schematic illustration showing that SpyCas9 mRNA and individual or group of three gRNAs were injected at one cell zebrafish embryo and phenotypes were observed at 6 hpf and 1 dpf as well as mutation rate was calculated by genotyping at 1dpf embryos. B. Boxplot is showing the high percentage of SpyCas9 indel mutation associated with zygotic copies of *cth1* gene at 1 dpf embryos from individual gRNAs as well as set of 3 combined gRNAs. Cas9 alone and *albino* gene are negative and positive controls respectively. n represents the numbers of embryos sequenced for each condition. C. Representative images and stack bar plot showing that no major developmental phenotypes were observed from F0 embryos injected with any of the guideRNAs and/or SpyCas9 combination. Different intensities of gray color are directly associated with severity of developmental phenotype (Class III (most severe) and white (wildtype). n represents the numbers of embryos analyzed at 1 dpf. Scale bar, 0.5mm. D. Representative images of adult *cth1* mutant fish (SpyCas9) with their corresponding wildtype controls. Top panel is for males and bottom for females. All the fish are 14 months old and apparently looking normal. Scale bar 1cm. E. Representative embryos images after crossing wildtype and/or SpyCas9 *cth1* F0 mutants (red color text) adults. Mutant females produced either dead or unfertilized embryos, mutants’ males produced unfertilized embryos, while wildtype crosses produced viable embryos. (Panels show 1.5 hpf, 5 hpf and 24 hpf embryos). Scale bar, 1000 μm. F. Representative images showing that oocytes from two independent *cth1* Cas9 mutant females are white, fragmented and degraded compared to oocytes from wildtype female. Scale bar 1000 μm. G. Representative scanning electron images of mature sperm from wild type and from independent *cth1* mutant males showing the abnormal morphology, wrinkled degraded head and tail with different morphologies ranging from no tail, short, to multiple fragmented tails phenotypes. Scale bar is 3 μm.

Although F0 mutants developed into apparently normal adults (Fig. 3D), both mutant males and females exhibited severe infertility problems (Fig. 3E). Crosses involving either *cth1* F0 mutant males or females with wildtype fish, or crosses between mutants, consistently resulted in unviable or degraded embryos (Fig. 3E). Adult F0 mutants generated with individual *cth1* gRNAs also show the similar results, ruling out non-specific gRNA effects (Fig. S3C). Further examination revealed substantial defects in gamete development. Mutant oocytes were translucent white and frequently ruptured, in stark contrast to the intact, pale-yellow wild-type oocytes (Fig. 3F). Similarly, sperm from *cth1* mutant males exhibited abnormal morphology, characterized by rough or degraded heads and various tail abnormalities (short, absent, bifurcated, or serrated tails), rendering them incapable of fertilizing wild-type eggs (Fig. 3G).

These results demonstrate that while zygotic *cth1* expression product is not critical for early embryogenesis, it plays a crucial role in fertility through the regulation of both oogenesis and spermatogenesis. The severe infertility phenotype highlights the necessity for alternative methods, such as conditional or temporal knockdown, to explore the functional role of *cth1* during early embryonic development.

### Cas13d-mediated knockdown reveals the essential role of *cth1* during the maternal-to-zygotic transition

Given that *cth1* germline mutants result in infertility and cannot be utilized to investigate early developmental roles (Fig. 3), we employed the RfxCas13d to transiently knock down *cth1* mRNA during embryogenesis (Hernandez-Huertas et al, 2022; Kushawah et al, 2020). Three independent gRNAs targeting *cth1* were designed and injected individually with Cas13d mRNA into one-cell stage embryos (Fig. 4A). The qRT-

**Figure 4.**
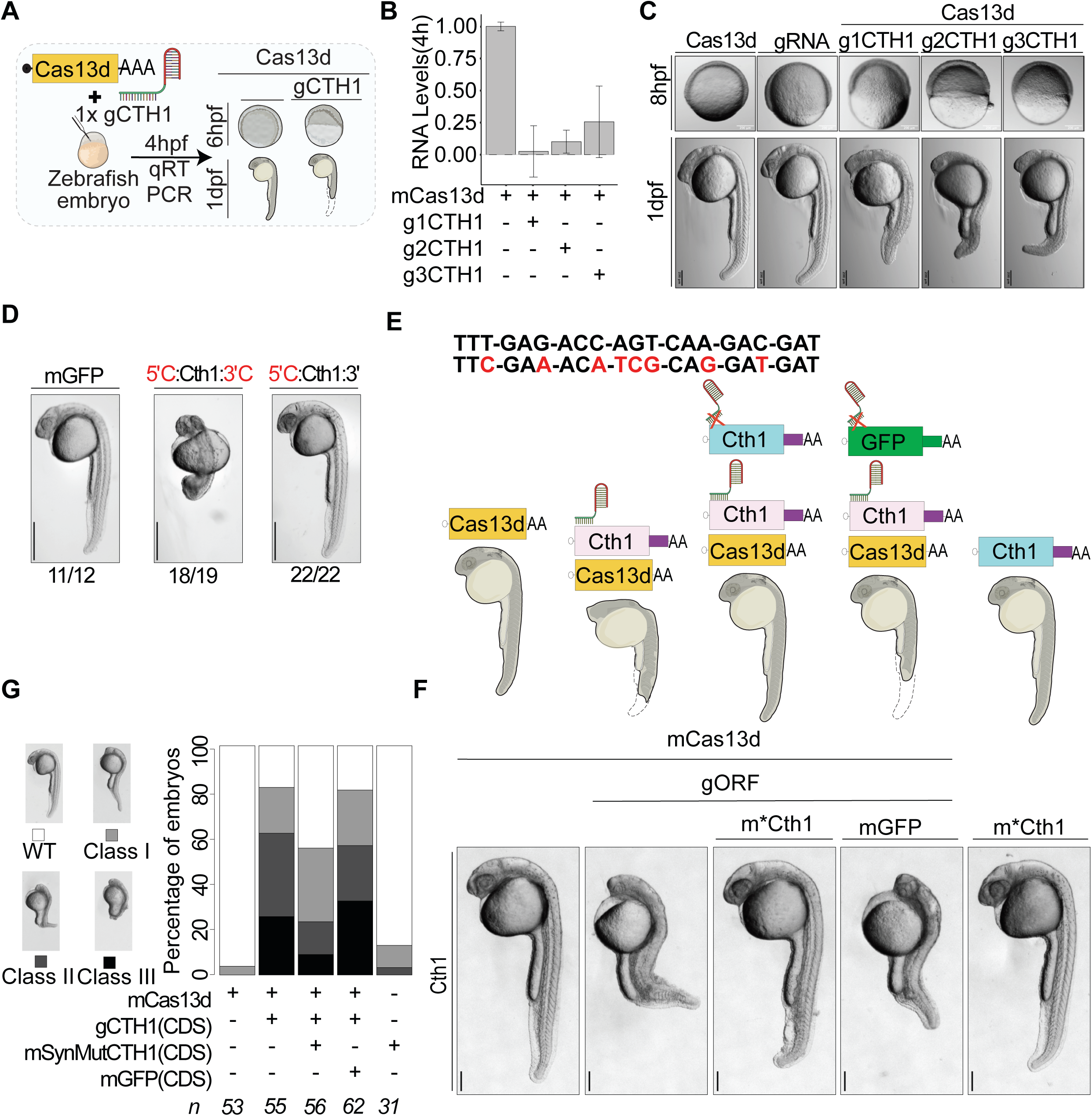
Cas13d-mediated knockdown reveals the essential role of *cth1* during the maternal-to-zygotic transition. A. Schematic illustration showing that RfxCas13d mRNA and/or three individual *cth1* gRNAs (g1CTH1, g2CTH1, g3CTH1) were injected into 1 cell stage embryo and qRT-PCR was performed at 4 hpf to test the target efficacy followed by imaging at 6 hpf and 1 dpf to observe embryonic development. B. Bar-plot showing *cth1* RNA levels upon CRISPR-RfxCas13d mediated knockdown for three independent gRNAs (g1CTH1, g2CTH1, g3CTH1) at 4 hpf measured by qRT-PCR. All the gRNAs show significant level of *cth1* mRNA knockdowns, (p-value ≤ 0.02 (g3CTH1), t. test). C. Representative images of embryos at 6 hpf and 1dpf. Embryos were co-injected with RfxCas13d and either of the guideRNAs targeting *cth1* displayed developmental abnormalities while injection of either RfxCas13d or guideRNA alone display normal embryonic development. Scale bar, 200 μm. D. Representative 1 dpf zebrafish images showing that injection of mRNA encoding for *cth1* with *control* 5’ and 3’UTRs affected development while injection of *cth1* mRNA with its endogenous 3’UTR does not affect development compared to embryos injected with mRNA encoding for GFP. Scale bar 5mm. E. Schematic illustration of RfxCas13d mediated RNA knockdown phenotype rescue assay in zebrafish. The CDS of endogenous *cth1* is targeted with a single g2CTH1 and RfxCas13d mRNA. For rescue experiment, ectopic *cth1* mRNAs containing 8 synonymous mutations in the gRNA target site (m*\*cth1(synMut)* RNA) and containing *cth1* 3’UTR should rescue the *cth1* knockdown phenotype. While ectopic expression of GFP should not rescue the developmental knockdown phenotype. F. Representative images showing the developmental phenotype associated with *cth1* knockdown (RfxCas13d + gORF), interestingly, the phenotype was rescued by ectopic expression of *cth1* insensitive to the guideRNA (RfxCas13d + gORF + *m*cth1(synMut)*) but not by the ectopic expression of GFP (RfxCas13d + gORF + mGFP). Injection of RfxCas13d alone or RfxCas13d + gORF and m*\*cth1(synMut)* mRNAs did not affect development. Images are of 28 hpf embryos, scale bar, 1mm. G. Stack bar plot showing the percentage of observed phenotype in the *cth1* rescue experiment at 1 dpf. Different intensities of gray color are positively associated with the severity of the phenotype (dark, most severe phenotype (class III) and white, (wildtype)). *n* represents the number of embryos observed in each condition.

PCR confirmed effective knockdown at 4 hpf, revealing significant reductions in *cth1* mRNA levels across all guides as compared to controls by at least 75% (p-value ≤ 0.02, Cas13d alone vs Cas13d and g3CTH1, t. test) (Fig. 4B). Phenotypic analyses demonstrated that embryos co-injected with Cas13d mRNA and any of the three individual gRNAs consistently exhibited developmental defects at both 6, 8 and 24 hpf (p-value ≤ 0.015, chi-square test) (Fig. 4C, S4A-C). These defects included delayed epiboly, abnormal axis elongation, and morphological abnormalities (Fig. 4C, S4A-C). Importantly, embryos injected only with Cas13d mRNA or with individual gRNAs alone did not display any of these developmental defects (Fig. 4C, S4A-C) (Hernandez-Huertas et al, 2022; Hernandez-Huertas et al, 2025; Kushawah et al, 2020; Moreno-Sanchez et al, 2025). The reproducibility of the developmental phenotypes across three independent gRNAs illustrates the specificity of the effects and minimizes the likelihood they arise through off-target effects. Collectively, these results indicate that maternal *cth1* is essential for embryogenesis (Figs. 4C, S4A-C), and comparison to *cth1* Cas9 mutants suggests that the early phenotype is primarily driven by maternally deposited *cth1* RNA rather than zygotic expression (Fig. S4D).

To further validate the functional specificity of the phenotypes, we performed rescue experiments. As a first step, we tested whether ectopic expression of *cth1* mRNA could perturb development. Injection of *cth1* fused to the control 3′UTR caused dose-dependent developmental defects and do not show marginal enrichment of RNA (Figs. 4D, S4E-G), whereas injection of *cth1* mRNA containing its native 3′UTR, which directs proper marginal localization (Fig. 2) did not show developmental defects (Fig. 4D and S4F). These findings highlight the importance of spatially regulated *cth1* expression during embryogenesis and demonstrate a need for preserving the native 3′UTR of *cth1* in rescue experiments.

Accordingly, we generated an exogenous *cth1* mRNA construct containing eight synonymous mutations resistant to a Cas13d gRNA (Fig. 4E). Co-injection of Cas13d, *cth1*-targeting gRNAs, and the resistant *cth1* construct harboring its native 3′UTR successfully rescued the knockdown developmental defects (p-value = 5.5e-4, Cas13d KD vs rescue 24 hpf, p-value = 2.4e-3, Cas13d KD vs rescue 6 hpf) (Figs. 4E-G, S4H - I). As control, GFP mRNA did not rescue the *cth1* phenotype and injection of *cth1* with its native 3′UTR alone did not affect development although both transcripts were efficiently translated (Figs. 4E-G, S4H-J). Moreover, *cth1* fused to a control 3′UTR failed to rescue the knockdown phenotypes even with minimal concentration (5pg/embryo) (Figs. S4K-M), highlighting the criticalness of spatial distribution of *Cth1* mRNA during gastrulation. Together, these results demonstrate that the phenotype observed upon *cth1* depletion are specific and biological relevant and that more generally Cas13d-mediated knockdown is an effective approach to study maternal genes essential for fertility and early development.

### *Cth1* regulates maternal mRNA stability during the maternal-to-zygotic transition independently of zygotic factors

Given that orthologs of Cth1 in other vertebrates’ function as RNA-binding proteins modulating mRNA stability (Ball et al, 2014; Cicchetto et al, 2023), we hypothesized that zebrafish CTH1 similarly regulates maternal mRNA clearance during the MZT. To test this, we performed RNA-seq on embryos subjected to *cth1* knockdown using Cas13d and two independent gRNAs (g1 and g2) at 4 hpf, a stage at which no evident developmental phenotypes were observed in *cth1* knockdown embryos (Fig. 5A). Both showed robust target efficiency (Fig. 4B), and reproducible phenotypes (Figs. 4C, S4A) that were rescued with exogenous *cth1* (Figs. 4E-G, S4H-I). RNA-seq profiles from embryos injected with *cth1*-targeting guides were compared to those from embryos injected with Cas13d alone at 2, and 4 hpf (Fig. 5B-C, S5A-B). Both gRNAs effectively reduced *cth1* mRNA levels across 2 and 4 hpf (2 hpf = log2 fold change < −0.9-fold reduction and adjusted p value ≤ 0.16, 4 hpf = log fold change < −1.95-fold reduction and adjusted p value ≤ 2.1 e-4) (Figs. 5B-C, S5A-B).

**Figure 5.**
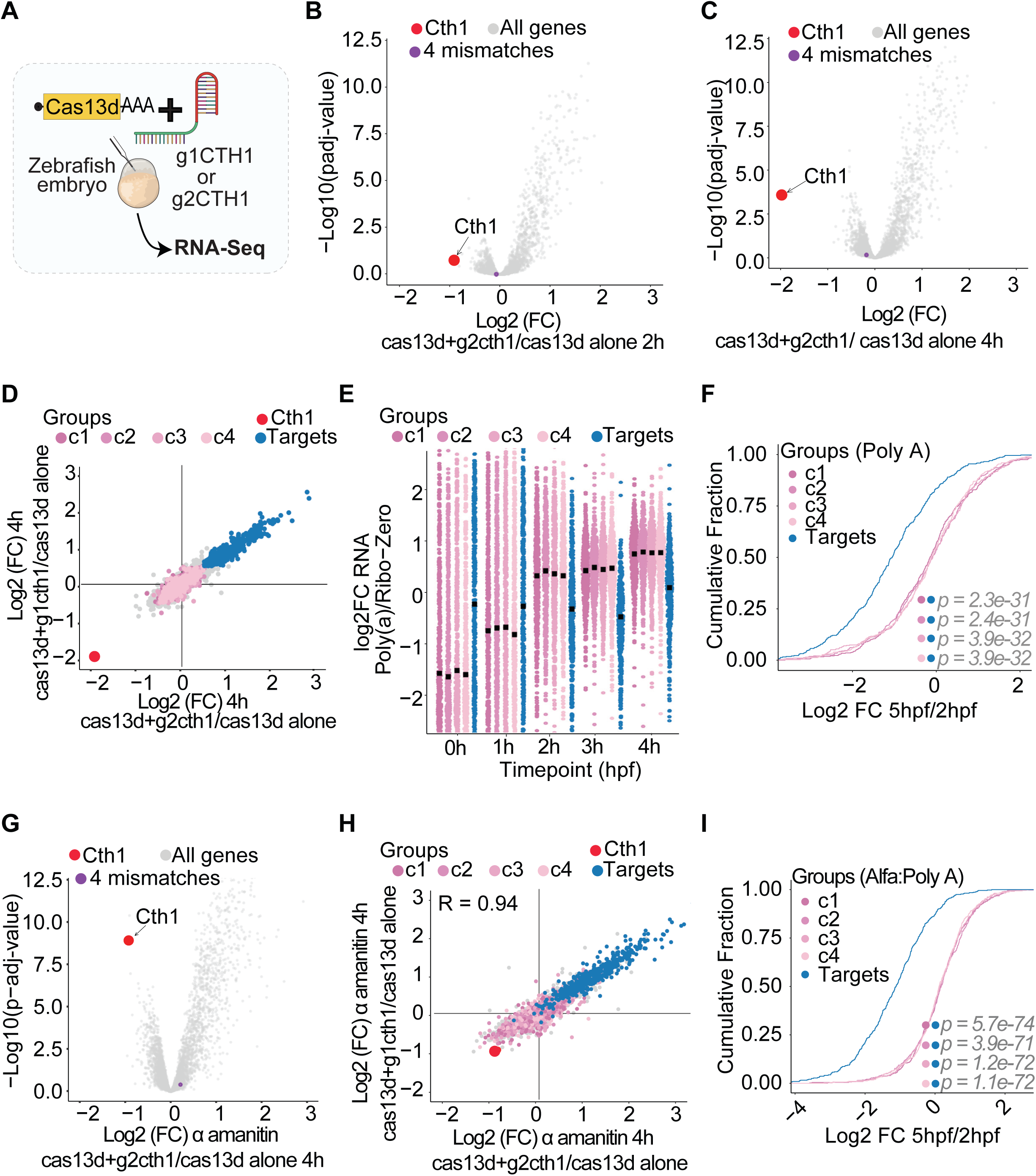
*Cth1* regulates mRNA stability during the MZT via 3′UTR *cis* elements. A. Schematic illustration showing that RfxCas13d mRNA was injected alone or with either gRNAs g1CTH1 or g2CTH1 at 1 cell stage zebrafish embryos and RNA-seq was performed at 2 and 4 hpi. B. Volcano plots showing the mRNA fold change in embryos co-injected with RfxCas13d and g2CTH1 compared to embryos injected with RfxCas13d alone at 2 (B) and 4 (C) hpf. *Cth1* was efficiently knockdown (red), and no off targets were identified allowing up to 4 mismatches between g2CTH1 sequence and the target sequences are shown by purple dots. D. Scatter plot showing the fold change of mRNA between the embryos co-injected with RfxCas13d and the indicated guideRNAs compared to the embryos injected with RfxCas13d alone. *Cth1* was effectively knockdown with both guideRNAs (Red), mRNAs significantly upregulated in both knockdown is shown in blue (putative Cth1 targets) and 4 random groups of 500 nonsignificant regulated gene as shown in pink color shades were selected as control genes (C1-C4). Strong positive correlation between the two-knockdowns ruling out off target effects (R = 0.95). E. Sina plot showing the relative poly(A) tail status, the log2 fold change between poly(A) RNA to ribosomal-depleted RNA, of the putative Cth1 targets and control groups (C1-C4) at different time points. Putative Cth1 targets exhibit extended poly(A) tail relative to control groups (C1-C4) immediately after fertilization (0 hpf) but progressively relative loss of poly(A) tail length over time during maternal zygotic transition. F. Cumulative plot showing the PolyA RNA-seq fold change between 5 and 2 hpf. The putative Cth1 targets (Blue) are unstable as compared to control groups (C1-C4, pink color). Wilcoxon rank-sum test, P value ≤ 2.3e-31. G. Volcano plots showing the mRNA fold change in α-amanitin treated embryos co-injected with RfxCas13d and g2CTH1 guideRNAs compared to embryos injected with RfxCas13d alone at 4 hpf. *Cth1* was efficiently knockdown (red), and no off targets were identified allowing up to 4 mismatches between g2CTH1 sequence and the target sequences are shown by purple dots which is suggesting *cth1* mRNA specific knockdowns even in α-amanitin treated embryos. H. Scatter plot showing the fold change of mRNA between the α-amanitin treated embryos injected with RfxCas13d and the indicated guideRNAs compared to the embryos injected with RfxCas13d alone. *Cth1* was effectively knockdown with both guideRNAs (Red), the defined Cth1 targets (D) were again upregulated (Blue) and the defined control groups not upregulated (pink shades). Strong positive correlation between the two-knockdowns ruling out off target effects (R = 0.94). I. Cumulative plot showing the PolyA RNA-seq plot fold change between 5 and 2 hpf from α-amanitin treated embryos. The putative Cth1 targets (Blue) are unstable as compared to control groups (C1-C4, pink color). Wilcoxon rank-sum test, P value ≤ 3.9e-71.

Knockdown of *cth1* led to more upregulated than downregulated genes for each gRNA at both 2 and 4 hpf (Fig. S5C). The sets of upregulated genes showed strong and significant overlap between the two gRNAs at each time point (2 hpf g1 vs. g2: p value ≤ 0.2e-16; 4 hpf g1 vs. g2: p value ≤ 0.2e-16; Fisher’s Exact Test) (Fig. S5C), as well as across developmental time points within each knockdown (2 vs. 4 hpf: p value ≤ 0.2e-16; Fisher’s Exact Test) (Fig. S5C). This reproducibility across guides and time points underscores the specificity of the observed transcriptional changes. Moreover, the fold change values strongly correlate between each gRNA knockdown and the control embryos (4 hpf = Pearson correlation R=0.95, p value ≤ 2.2e-16) (Fig. 5D), suggesting that these genes are direct or indirect *cth1* targets rather than off-target effects (Fig. 5D). This is also supported by our off-target analysis, where down regulated genes did not align to either gRNAs allowing up to 4 mismatches (shown in purple) (Figs. 5B-C, S5A-B). The significant overlap of upregulated mRNAs at 2 and 4 hpf (Fig. S5C), together with the observation that many changes were already evident by 2 hpf, indicates that *cth1* exerts a rapid maternal effect.

To define high confidence potential *cth1* targets, we selected genes upregulated in both knockdown conditions (fold change ≥ 0.5 and p value ≤ 0.05, putative *cth1* targets (Targets)) and as controls, we generated four randomly sampled groups of 500 genes (C1-C4, shown in different shades of pink) from the transcriptome that were not significantly changed with either of the gRNAs targeting *cth1* (Fig. 5D). High-confidence targets group (Targets) displayed elevated poly(A)-tail status at 0 hpf relative to control groups, suggesting these mRNAs are maternally deposited with extended poly(A) tails (Fig. 5E). During the MZT, these mRNAs (Targets) underwent progressive poly(A) shortening (Fig. 5E) and reduced expression compared to controls (C1-C4) at respective time points (Fig. 5F and S5D-E), indicative of active deadenylation and degradation processes.

Since *cth1* is maternally supplied (Fig. 1), its decay is zygotic transcription-independent (Fig. 1D) and the observation that targets were affected before zygotic genome activation (Figs. 5B, S5A) suggests that *cth1* mediated mRNA decay is governed solely by maternal factors (maternal program). To validate this, we repeated *cth1* knockdown using Cas13d in embryos co-injected with α-amanitin to block zygotic transcription. *cth1* knockdown was effective under these conditions (fold change ≥ 0.5 and p value ≤ 0.05) (Fig. 5G, S5F) and potential *cth1* targets continued to display significant upregulation compared to controls (Fig. 5H). These results confirm that *cth1* mediated mRNA degradation is independent of zygotic transcription. Further analysis of RNA-seq time-courses from embryos injected with α-amanitin revealed a consistent and progressive decay of *cth1* target mRNAs, reinforcing the conclusion that their regulation is maternally governed and independent of zygotic transcriptional activity (Figs. 5I and S5G).

This application of CRISPR-Cas13d combined with α-amanitin treatment provides a unique method for accurately assessing maternal mRNA stability independent of zygotic transcriptional activity. Collectively, our findings strongly support a model wherein *cth1* functions within a purely maternal regulatory program, directly or indirectly governing maternal mRNA stability during the MZT.

### *Cth1* regulates mRNA stability during the MZT via 3′UTR *cis* elements

To determine whether *cth1* regulates its targets through elements within their 3′UTRs, we searched for *cis*-regulatory motifs enriched in these regions. As candidate targets, we again used our selected genes (Fig. 5D) that were significantly upregulated by both individual *cth1*-targeting gRNAs at 4 hpf compared to embryos injected with Cas13d alone. As controls, we also used our four independent sets of 500 genes randomly sampled mRNAs which were not significantly upregulated in either knockdown condition (Fig. 5D). Candidate targets did not differ significantly from controls in 3′UTR lengths (p value ≥ 0.56, Fig. S6A), ruling out sequence length as a confounding factor for motif analysis.

Among the top enriched 8-mers and 7-mers in the 3′UTR of the potential *cth1* targets compared to the control groups (Figs. 6A and S6B), the 8-mers AAUAAAUA was present in 131 (29%) and AAUAAUAA in 66 (15%) of the 444 candidate target genes (Fig. 6B).

**Figure 6.**
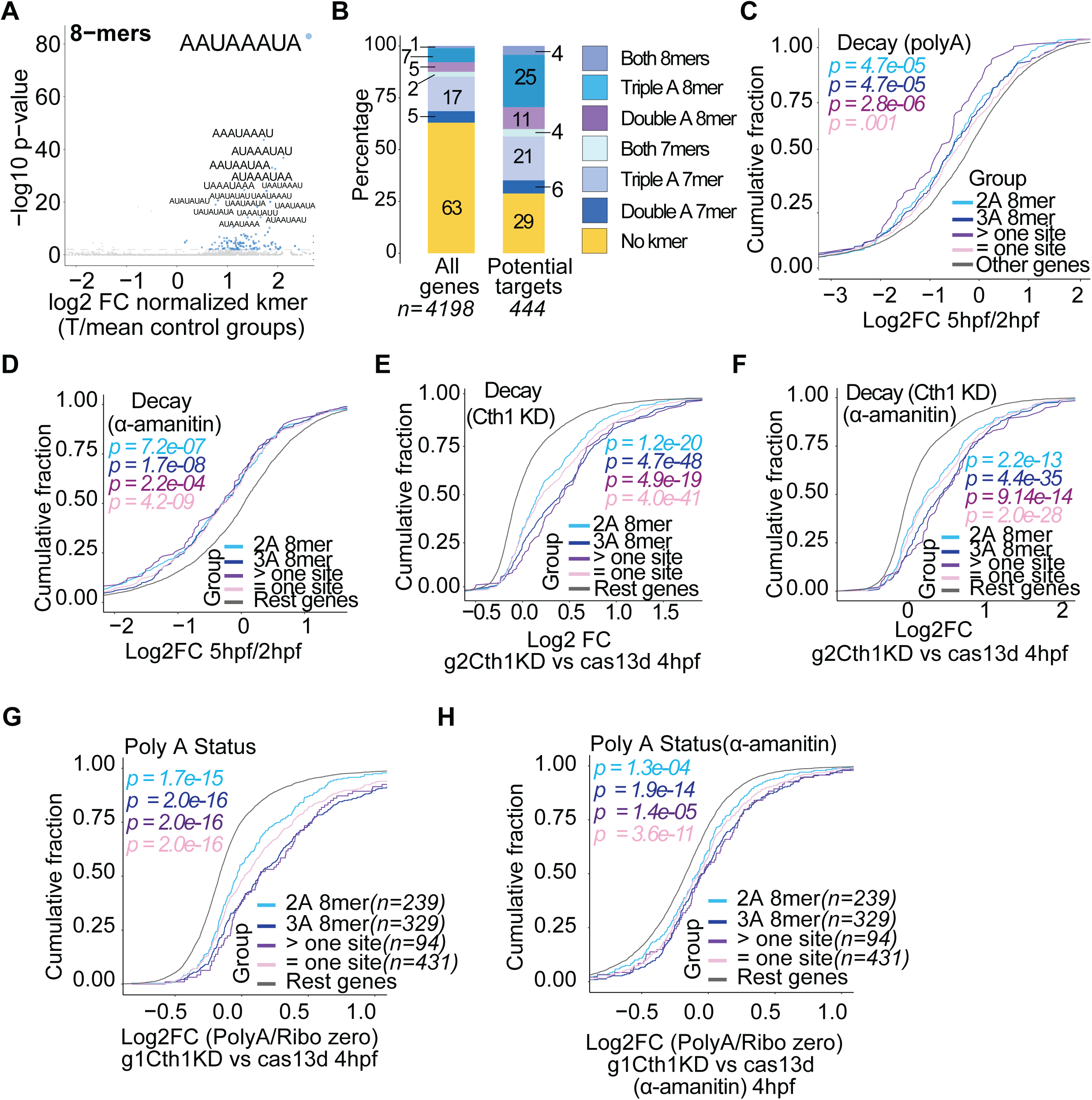
Cth1 acts on the AU - rich 3′UTR of its targets. A. Scatter plot showing 8-mer motifs that are enriched in the putative Cth1 target group compared to the 4 control groups (C1-C4). AU-rich motifs are particularly overrepresented, with sequences containing “AAA” and “AA” showing the strongest enrichment. B. Percentage of the 7-mer and 8-mer motifs containing “AAA” and “AA” one or more times in the putative Cth1 target and all the genes. The putative Cth1 target group genes with enriched motifs with these combinations constitute more than 70% of total genes, while in all the genes constitution is 37%. C. Cumulative plot showing fold change between PolyA RNA-seq at 5 and 2 hpf. Transcripts containing enriched 7-mers/8-mers, particularly those with “AAA” and multiple motifs, exhibit greater destabilization than other genes. Wilcoxon rank-sum test, P values indicated. D. Cumulative plot showing fold change between PolyA RNA-seq plot from α-amanitin treated embryos at 5 and 2 hpf. Transcripts containing enriched 7-mers/8-mers, especially those with “AAA” and multiple motifs, exhibit greater instability than other genes. Wilcoxon rank-sum test, P values indicated. E. Cumulative plot showing the fold change between in PolyA RNA-seq upon *cth1* knockdown mediated by g2CTH1 at 4 and 2 hpf. Transcripts containing enriched 7-mers/8-mers, especially those with “AAA” and multiple motifs, exhibit more stability than other genes. Wilcoxon rank-sum test, P values indicated. F. Cumulative plot showing the fold change between fraction in PolyA RNA-seq plot upon *cth1* knockdown mediated by g2CTH1 in α-amanitin treated embryos at 4 and 2 hpf. Transcripts with “AAA” and multiple motifs remain more stable than other genes. Wilcoxon rank-sum test, P values indicated. G. Cumulative distribution plot of poly(A) status, as the log2 fold change between poly(A)-selected and ribo-depleted RNA-seq, upon *cth1* knockdown by g1Cth1KD versus Cas13d control at 4 hpf. Transcripts containing enriched 7-mers/8-mers, especially those with “AAA” and multiple motifs, show significantly greater poly(A) tail enrichment relative to other genes. Wilcoxon rank sum test, number of genes (*n*) and *p*-values are indicated. H. As in G, but in α-amanitin treated embryos using g1Cth1KD targeting *cth1*, confirming the reproducibility of the poly(A) tail enrichment observed for 7-mers/8-mers - containing transcripts upon *cth1* knockdown at 4 hpf.

The respective 7-mers, AUAAAUA and AUAAUAA, were found in 110 (25%) and 44 (10%) of the same group (Fig. 6B). Altogether, more than 71% of candidate targets contained at least one of these two seed motifs (AUAAAUA and AUAAUAA), whereas fewer than 37% of all analyzed genes carried any of them, representing nearly a two-fold enrichment (Fig. 6B).

The AAUAAAUA motif showed a strong 3′-biased distribution within the 3′UTRs (P = 3.3e-3 (non-targets), Wilcoxon rank-sum test), with significant enrichment in the final 30 nucleotides (P = 7.8e-4; Wilcoxon rank-sum test) (Fig. S6C). In contrast, AAUAAUAA did not display a positionally biased distribution (Fig. S6C). RBP-scan (Kretov et al, 2026), a recently developed *in vivo* RNA-binding activity tool, independently identified AU-rich motifs interactions upon *cth1* overexpression, supporting the presence of functional *cis*-elements recognized by Cth1 (Fig. S6D).

To evaluate whether the presence of these motifs was associated with mRNA instability during embryogenesis, we examined maternal mRNAs containing AAUAAAUA or AAUAAUAA motifs in their 3′UTRs. These genes displayed significantly greater degradation (5 vs. 2 hpf fold change) compared to mRNAs without these motifs, in both untreated (Figs. 6C, S6E-F) and α-amanitin–injected embryos (Figs. 6D, S6G-H). This suggests that motif-containing mRNAs are intrinsically unstable and degraded independent of zygotic transcription. Furthermore, these mRNAs containing the motifs were significantly upregulated in *cth1* knockdowns (Figs. 6E-F, S6I-J), indicating they are *cth1* targets (direct or indirect). 3′UTRs containing more than one motif showed stronger decay than those with a single motif during embryogenesis (Fig. 6C, S6E-F) and higher stabilization in *cth1* knockdown (Fig. 6E-F, S6I-J), suggesting that stability of the 3′UTR is directly associated with motif numbers. In both untreated and α-amanitin–injected embryos, *cth1* knockdown led to higher poly(A)-tail status on *cth1* target mRNAs, with tail length scaling positively with both the number and length of motifs (Fig. 6G-H, S6K-L). This is consistent with the progressive poly(A) tail shortening of these transcripts observed during embryogenesis (Fig. 5E), a process that proceeds independently of zygotic transcription (Fig. S5D), and indicates that Cth1 is required for their deadenylation-mediated decay (Fig. 6G, 6H, S6K, S6L).

### Cth1 recognizes conserved AU-rich motifs in target 3′UTRs to drive deadenylation-mediated decay

To directly test whether *Cth1*-mediated destabilization depends on the motifs we identified as enriched in its target 3′UTRs (Fig. 6), we cloned 3′UTRs from representative targets containing at least one AAUAAAUA site (*irf5, lipia, ppdpfb, stx11a,* and *tmem243a*) downstream of GFP. These were co-injected with a GFP-control 3′UTR and *NanoLuc* as internal controls into Cas13d-alone (control) and *cth1* knockdown embryos (Cas13d and g1CTH1) (Fig. 7A). The qRT-PCR confirmed efficient *cth1* knockdown and, interestingly, most GFP-3′UTR reporters containing the motifs were significantly stabilized in *cth1* knockdown embryos at 6 hpf (p value ≤ 0.07, t. test) (Fig. 7B). Furthermore, the *tmem243a* 3′UTR, which contains two such motifs, showed the strongest effect (Fig. 7B) (p value ≤ 3.9e-07, t. test), reinforcing the influence of motif numbers (Fig. 6). The GFP-control 3′UTR control was unaffected (p value = 0.11, t. test), confirming specificity (Fig. 7B). Individual injection of GFP-3′UTR reporters containing these motifs also show higher GFP fluorescence levels upon *cth1* knockdown as compared to wildtype embryos at 8 hpf (Figs. 7C, S7A-C) All these results suggest that *cth1* promotes their decay via 3′UTR interactions.

**Figure 7.**
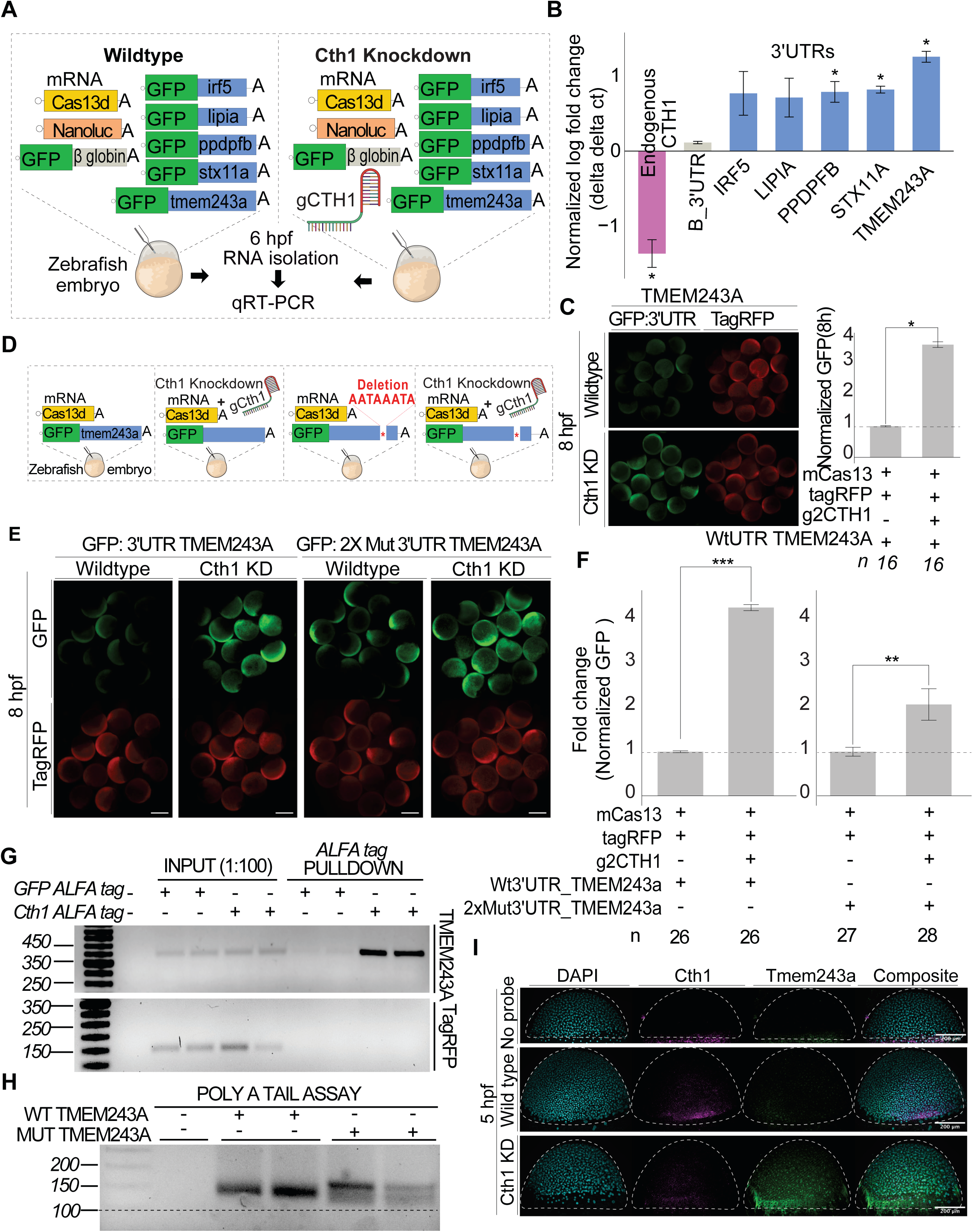
Cth1 recognizes conserved AU-rich motifs in target 3′UTRs to drive deadenylation-mediated decay. A. Schematic of the GFP:3′UTR reporter assay in wild-type and RfxCas13d-mediated *cth1* knockdown conditions followed by qRT-PCR at 6hpi. 3′UTRs of five candidates from putative Cth1 target group were cloned downstream of GFP, with injection controls including GFP: *β-globin* 3′UTR and NanoLuc mRNA (10 pg/embryo). This injection cocktail was injected into one-cell zebrafish embryos with or without g1CTH1 (600 pg/embryo) targeting *cth1*. B. qRT-PCR analysis showing normalized RNA levels in embryos injected with a cocktail of GFP:3′UTRs mRNAs, NanoLuc as control and RfxCas13d, with or without g1CTH1 at 6 hpf. Endogenous *cth1* transcripts were reduced while GFP: *β-globin* 3′UTR control remained unchanged and GFP:3′UTRs of all candidate targets were upregulated upon *cth1* knockdown. Data represent ≥3 biological replicates from two independent experiments; significance was determined by *t*-test (*p* < 0.05). C. Pictures showing the GFP expression from a mRNA encoding for GFP containing the 3′UTR for *tmem243a* reporter co-injected with RfxCas13d and g1CTH1 (KD) or in wildtype conditions, at 8 hpi. TagRFP was co-injected as a control. Bar plot showing the normalized GFP quantification in wildtype and *cth1* knockdown conditions. “n” number of embryo calculated, and significance was determined by *t*-test (*p* < 0.05). D. Schematic of the GFP:3′UTR reporter assay comparing reporters containing 3′UTR with intact endogenous motifs versus motif-deleted versions in wild-type and RfxCas13d-mediated *cth1* knockdown embryos at 8 hpf. Each reporter was injected individually along with TagRFP as injection control at the one-cell stage. E. Pictures showing the GFP expression from a mRNAs encoding for GFP containing the 3′UTR for *tmem243a (wildtype)* or lacking the proposed CTH1 motif reporters co-injected with RfxCas13d and g1CTH1 (KD) or in wildtype conditions, at 8 hpi. TagRFP was co-injected as a control. F. Bar plot showing the normalized GFP quantification for 3′UTR for *tmem243a (wildtype)* and lacking the proposed CTH1 motif reporters in wildtype and *cth1* knockdown conditions. “n” number of embryo calculated, and significance was determined by *t*-test (*p* < 0.05, *p* < 0.01). G. ALFA tag RNA immunoprecipitation (RIP) followed by RT-PCR to assess binding of Cth1 to the *tmem243a* 3′UTR. Input samples (1:100 dilution) confirm the presence of RNA across all conditions. Strong and specific enrichment of the *tmem243a* 3′UTR is detected in pull-downs from Cth1–ALFA tag–injected embryos, but not from GFP–ALFA tag control embryos, suggesting direct association of Cth1 with this target mRNA. Co-injected *tagRFP* mRNA serves as an injection and specificity control; it is detectable in input fractions but absent from pull-down fractions, confirming the selectivity of the enrichment. H. RT-PCR analysis of poly(A)-tail length assay for GFP reporters containing either the wild-type *TMEM243A* 3′UTR or a mutant version lacking the Cth1 binding motifs at 4 hpi. The wild-type reporter shows a shorter PCR product than the mutant, consistent with Cth1-dependent deadenylation. NT: No template control; WT: wild-type 3′UTR reporter; Mut: mutant 3′UTR reporter lacking Cth1 binding motifs. I. HCR analysis of endogenous *cth1* and *tmem243a* mRNA expression in wild-type and *cth1* knock-down (Cth1 KD) zebrafish embryos at 5 hpf. In wild-type embryos, *cth1* mRNA (Magenta) is enriched at the embryonic margin, while *tmem243a* mRNA (Green) is lowly expressed. Upon *cth1* knockdown, *tmem243a* mRNA (Green) exhibits markedly enhanced marginal localization, suggesting that Cth1 restricts the marginal enrichment of *tmem243a* transcripts. No-probe embryos were included as negative controls to confirm signal specificity. Cyan: DAPI (Nuclei), rightmost panel: Composite images, Scale bar, 200 μm.

**Figure 8.**
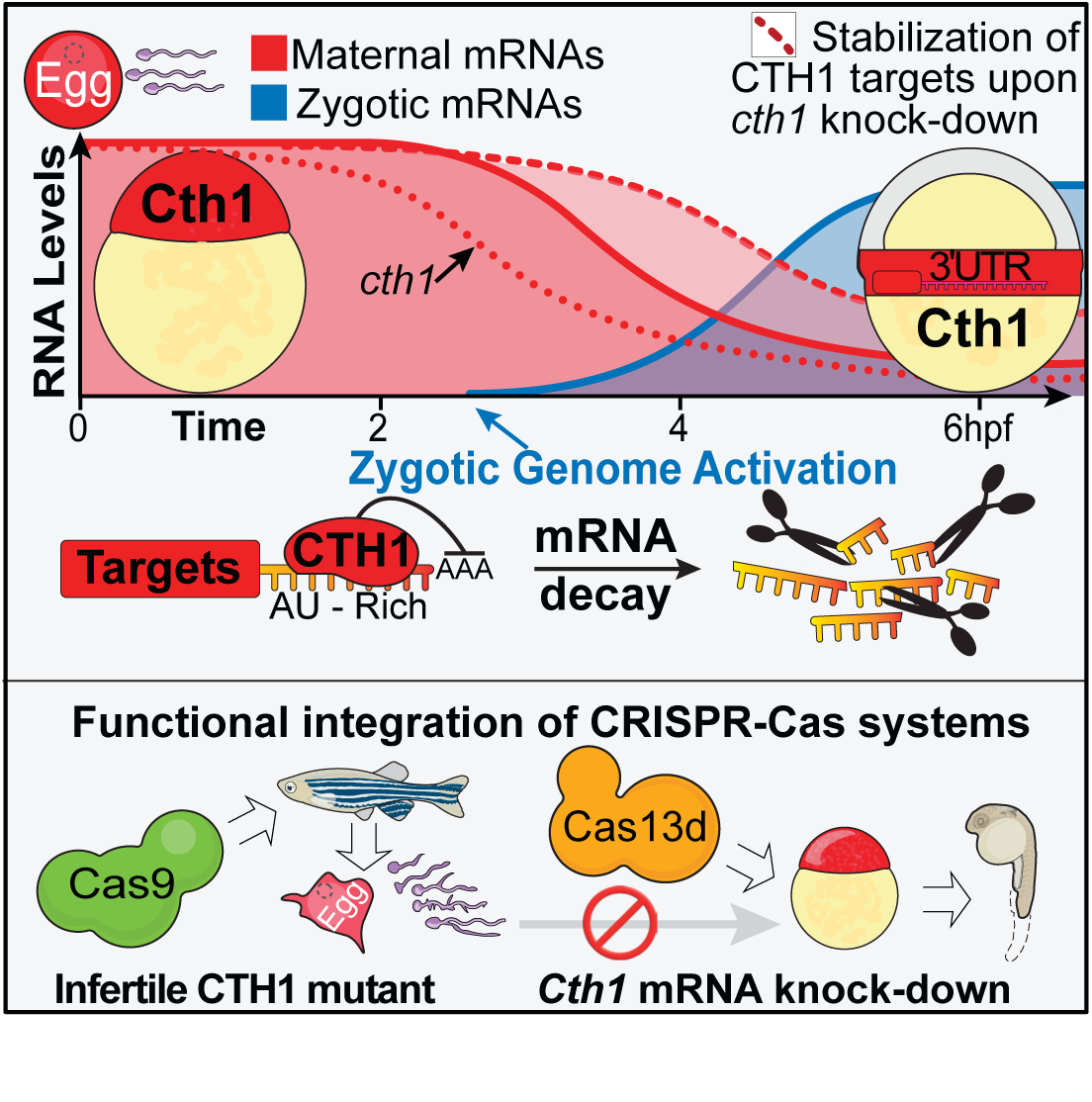
Functional analysis of *cth1* during maternal-to-zygotic transition. A. *Cth1* is among the most highly deposited maternal mRNAs and undergoes rapid, cell–specific degradation during the MZT. The *cth1* 3′UTR mediates mRNA stabilization in the marginal region of zebrafish embryos by 6 hpf. Knockdown of *cth1* using CRISPR–Cas13d affects embryonic development and stabilizes maternal mRNA targets which have AU-rich 3′UTRs by preventing their deadenylation. In contrast, CRISPR–Cas9-generated *cth1* mutants are infertile, underscoring an essential role for *cth1* in both oogenesis and spermatogenesis. The Cas13d approach enabled functional interrogation of *cth1* during embryogenesis, bypassing the embryonic lethality observed in Cas9 mutants.

To assess the requirement of these motifs within the 3′UTRs, we generated deletion constructs lacking AAUAAAUA motifs for *tmem243a* 3′UTR. These mutant reporters were individually co-injected with TagRFP as an internal control into Cas13d-alone and *cth1* knockdown embryos (Fig. 7E). In all cases, GFP expression from mutant-motifs (deleted version) 3′UTRs was increased compared to endogenous versions (wildtype), (Fig. 7E-F), confirming that motif and *cth1* both are required for mRNA destabilization.

To assess whether Cth1 physically interact with its targets, we performed RNA immunoprecipitation followed by RT-PCR (RIP–RT-PCR) using an antibody against tagged Cth1. This confirmed a direct interaction between Cth1 and *tmem243a* mRNA (Fig. 7G). To then test whether this interaction drives deadenylation through the identified motifs, we performed poly(A) tail assays on wild-type and motif-deleted *tmem243a* 3′UTR reporters. These experiments revealed that Cth1-mediated poly(A) shortening is motif-dependent (Figs. 7H, S7D), directly linking motif recognition to the deadenylation. Finally, HCR analysis in *cth1* knockdown embryos revealed preferential accumulation of *tmem243a* and *lipia* mRNAs at the embryonic margin during gastrulation (Figs. 7I, S7E), indicating that Cth1 accumulation in the marginal (Fig. 2) prevent the accumulation of these transcripts in this region. In summary, our results identify Cth1 as a key maternal decay factor that drives deadenylation-mediated clearance of its targets through conserved cis-regulatory elements in their 3′UTRs, thereby establishing the spatiotemporal pattern of maternal mRNA decay during embryogenesis.

## Discussion

Our study demonstrates that differential gene expression during early development can result not only from transcriptional activity but also from spatially and temporally regulated mRNA decay. *Cth1* emerges as the first maternal RNA decay factor in zebrafish with spatially restricted accumulation outside of germline contexts. Its high maternal abundance, rapid degradation, and 3′UTR-dependent marginal localization (Figs. 1, 2 and 7) underscore a novel layer of post-transcriptional control shaping the embryonic transcriptome.

The marginal enrichment of *cth1* is most consistent with spatially regulated degradation rather than localized transcription or directed mRNA transport. Injection of reporter mRNAs containing the *cth1* 3′UTR confirmed that this localization is encoded within the 3′UTR sequence (Fig. 2). Interestingly, the GFP signal was also mainly coming from the marginal region, suggesting that the endogenous Cth1 protein might also accumulate and function in the marginal region (Fig. 2C), which is supported with the spatio-temporal mis regulation of the Cth1 targets (Fig. 7I). Massively parallel reporter analysis of the *cth1* 3′UTR identified three unstable regions that localize the mRNA to marginal region, suggesting that overlapping cis-regulatory elements and/or region-specific trans-acting factors may protect these sequences from decay (Fig. 2). The evolutionary conservation of these motifs (Fig. 2H) suggests selective pressure to preserve this mechanism and a valuable entry point for identifying analogous regulatory elements in other maternal mRNAs in the future. Our findings challenge aspects of traditional transcriptional models by demonstrating that vertebrate maternal RNAs can be spatially patterned through differential degradation outside the germ layer context. In *Drosophila* germ cells, Oskar protects *nanos* mRNA from Smaug-mediated deadenylation, allowing spatially restricted *nanos* mRNA stabilization and translation(Pamula & Lehmann, 2024; Semotok et al, 2005; Siddiqui et al, 2024; Tadros et al, 2007; Zaessinger et al, 2006). However, such *trans*-acting factors mediating *cth1* spatio-temporal regulation remain unknown. A direct or indirect autoregulatory role of Cth1 can be part of the mechanism as shown for its orthologous in yeast (Martinez-Pastor et al, 2013). Identifying RNA-binding proteins or cofactors that interact with the *cth1* 3′UTR triggering its decay and/or protecting it from degradation may reveal additional components of a broader spatiotemporal decay network.

The discovery of *cth1* decay-mediated localization raises the possibility that additional mRNAs may be spatio-temporally localized at the marginal region of the embryo or in other regions through selective degradation. The high abundance of *cth1* may have facilitated its identification, suggesting it is merely the visible tip of a broader regulatory network. Future spatially resolved approaches, such as single-cell RNA-seq, whole-embryos imaging platform, SLAM-seq, and injection of massive reporter libraries will be instrumental in uncovering additional maternal mRNAs regulated by spatially restricted decay (Bhat et al, 2023; Fishman et al, 2024; Rabani et al, 2018). These technologies offer single-cell and transcript-level resolution, enabling the discovery of new maternal regulators. As shown recently, using single cell SLAM-seq in zebrafish, different mRNAs follow differential cell specific decay rate and patterns associated with specific sequence elements (Fishman et al, 2024). This study also identified *cth1* as a rapidly decaying maternal RNA and suggested that zygotic *cth1* transcripts may contribute to development (Fishman et al, 2024). However, our data support a predominant maternal role for *cth1* during embryogenesis (Figs. 3 and 4) but does not exclude a potential zygotic contribution, which will require further detailed investigation.

Here, we applied for the first time a dual CRISPR strategy, Cas9 and Cas13d, to dissect maternal versus zygotic gene functions during vertebrate embryogenesis. Cas9-based mutagenesis did not result in early developmental defects (Fig. 3), suggesting that zygotic *cth1* is dispensable at early developmental stages. Although *cth1* mutants generated by Cas9 apparently develop normally but exhibit severe fertility defects later in life due to impaired gametogenesis (Figs. 3 and7). Similar phenotypes have been reported for *Cth1* orthologs in other species (Ball et al, 2014; Ramos, 2012; Zhou et al, 2023) which hinders the functional investigation of these orthologs during early embryonic development (Ramos et al, 2004; Sha et al, 2018), though these have been implicated in female fertility. Our results suggest that *Cth1* function extends to both male and female gametogenesis in zebrafish (Figs. 3 and 7). Detailed analysis of these mutant fish revealed that Cth1 is required for early meiotic progression and is essential for gametogenesis and fertility in both male and female zebrafish (Kushawah et al, 2026).

This positions zebrafish as a powerful model to investigate *Cth1*-related infertility and other potential developmental roles. Although this infertility phenotype prevents the use of loss-of-function mutants to assess maternal function during embryogenesis. This limitation was overcome by using Cas13d-mediated knockdown, which enabled targeted depletion of maternal mRNAs and assessment of their developmental roles (Figs. 4, 5, 6 and 7). Moreover, we combined Cas13d with α-amanitin treatment, providing a new approach to test maternal mRNA stability independent of zygotic transcription. Furthermore, while collateral activity has been reported for Cas13d in mammalian systems (Ai et al, 2022; Kelley et al, 2022; Li et al, 2023; Shi et al, 2023), there is no evidence to suggest this occurs in zebrafish embryos injected with RfxCas13d protein or mRNA and gRNA targeting endogenous mRNAs (Cheng et al, 2022; Hernandez-Huertas et al, 2025; Kushawah et al, 2020; Moreno-Sanchez et al, 2025; Shangguan et al, 2024; Treichel & Bazzini, 2022; Treichel et al, 2026; Zhou et al, 2020). The use of two independent gRNAs (Fig. 4), rescue of the phenotypes by ectopic expression of *cth1* (Fig. 4), and the overlap of differentially expressed mRNAs across gRNAs (Fig. 5) collectively confirm the specificity of the Cas13d approach used in our experiments (Fig. 5). The different developmental outcomes observed with ectopic *cth1* expression from heterologous (*control*) versus endogenous 3′UTRs (Fig. 4) underscore the importance of spatial regulation of decay factors themselves during embryogenesis. Altogether, the spatiotemporal regulation of *cth1*, its marginal localization, mRNA decay activity, and associated phenotypes, suggests that *Cth1* may influence mesodermal patterning and regulation of cells/tissues derived from mesodermal cell lineage differentiation during development. Defining how this mode of regulation shapes lineage specification, cell-fate decisions, and proper morphogenesis during gastrulation will be an important direction for future work.

Our knockdown experiments revealed widespread transcript upregulation detectable before ZGA as well as in α-amanitin–treated embryos, establishing that *Cth1* operates within the maternal program, independent of zygotic expression (Figs. 1 and 5). *Cth1* is a maternal RNA decay factor part of the maternal program in zebrafish. Mechanistically, *Cth1* promotes deadenylation and decay analogous to as other factors (Cicchetto et al, 2023; Lai et al, 2003), by recognizing AU-rich motifs (8-mers and 7-mers) in target 3′UTRs (Fig. 6 and 7). One of these motifs partially overlap with the poly(A)-tail signal (Gallicchio et al, 2023; Levitt et al, 1989; Ulitsky et al, 2012). mRNAs bearing these motifs are maternally deposited with extended poly(A) tails (Fig. 5) yet are preferentially destabilized during the MZT (Figs. 5, 6 and 7), with decay strength increasing with motif copy number (Figs. 6 - 7). AU-rich binding is independently supported by high-throughput reporter assays (RBP-scan; (Kretov et al, 2026)), prior iCLIP data showing CTH1 preference for 3′UTRs (Vejnar et al, 2019) and our RIP–RT-PCR demonstrating motif-dependent pulldown of target mRNA by (Fig. 7G). Deleting these motifs in 3′UTR reporters attenuated its deadenylation, decay as well as response to *cth1*, confirming their necessity for *Cth1*-dependent destabilization (Fig. 7). Consistent with this, massively parallel reporter works have previously linked AU-enriched sequences to mRNA instability during the MZT (Rabani et al, 2018; Vejnar et al, 2019). Finally, in *cth1* knockdown embryos we observed accumulation of Cth1 targets at the embryonic margin, indicating that marginal Cth1 enrichment (Fig. 2) normally restricts these transcripts from this region. The ectopic reporters did not display the same spatial pattern, which we hypothesize may reflect saturation of the Cth1-mediated decay system by the injected mRNA, or insufficient spatial resolution of the reporter assay to detect subtle marginal differences. Our observation of Cth1 enrichment in the marginal region raises the possibility that Cth1 protein levels may be controlled by spatially restricted mechanisms. In *C. elegans*, a conceptually similar mechanism regulates the stability of CCCH zinc-finger proteins during early embryogenesis (DeRenzo et al, 2003; Gallo et al, 2008; Oldenbroek et al, 2012; Schubert et al, 2000). The germline determinant PIE-1 is degraded in somatic blastomeres through ZIF-1-mediated ubiquitin-dependent proteolysis, while remaining stable in the germline lineage (DeRenzo et al, 2003; Oldenbroek et al, 2012; Schwartz et al, 2023). This spatially restricted turnover contributes to the segregation of somatic and germline fates. Although PIE-1 and Cth1 may not be direct functional equivalents, this example illustrates how zinc-finger proteins can be regulated at the level of protein stability in a cell-type-specific manner. By analogy, Cth1 localization in the marginal region may reflect a regulated stability mechanism, although this possibility requires direct experimental validation (DeRenzo et al, 2003; Oldenbroek et al, 2012; Schwartz et al, 2023).

This work opens new areas of investigation. (1) How many additional genes are regulated in a spatiotemporal manner through differential RNA stability (Kushawah et al, 2024)? (2) Which *trans*-acting factors recognize *cis*-elements and mediate localized decay of *cth1*? (3) How is *Cth1* protein distributed across the embryo, and more broadly, might its activity, rather than abundance, be controlled by spatially restricted post-translational modifications?

In summary, this study identifies *Cth1* as an RNA decay factor in the maternal program of zebrafish embryogenesis and first mRNA shown to be spatio-temporally distributed outside the germline. Our findings provide the first direct evidence that spatio-temporally regulated mRNA decay, outside the germline’s context, contributes to early development regulation during the MZT, establishing that RNA clearance, like transcription, can determine mRNA distribution in both spatial and temporal dimensions. While such regulatory logic likely operates during other cellular transitions, its effects may be technically easier to identify during early embryogenesis, when the transcriptome is dominated by maternally deposited mRNAs and different cell lineages are being established during gastrulation. These insights redefine our understanding of spatial gene regulation and offer a powerful conceptual and experimental framework for dissecting post-transcriptional gene regulation *in vivo* in a spatio-temporal manner.

## Methods

### Zebrafish maintenance, embryo productions and experimentations

All zebrafish (*Danio rerio*) experiments were performed in accordance with protocols approved by the Institutional Animal Care and Use Committee (IACUC) at the Stowers Institute. Adult zebrafish from the AB, TU, TF, and TLF strains, aged between 6 to 18 months, were used for embryo production. Fish were randomly selected from a colony of approximately 500 individuals, consisting of four independent strains. For microinjections, at least 12 male and 12 female zebrafish were randomly bred to get fertilized timed one cell stage embryos. Embryos were maintained in zebrafish embryo media at 28.5°C under standard laboratory conditions.

### Zebrafish microinjections

All the experiments were done by injecting one cell stage dechorionated zebrafish embryos. All the injected controls and experiment embryos were grown at 28.5°C using E2 zebrafish embryo media with 0.01% methylene blue. For RfxCas13d-mediated knockdown experiments, 300 pg of RfxCas13d mRNA (Addgene plasmid # 141320) was injected per embryo (Kushawah et al, 2020). Guide RNA (gRNA) concentrations for Cas13d knockdowns ranged from 400-600 pg. *Cth1* gRNAs were ordered from synthego. For Cas9 mutagenesis, 100pg of Cas9 mRNA (Addgene plasmid # 46757) and 25pg of each gRNA were used per injection. All the injections were done at one cell stage embryo. Following microinjections, all embryos were treated in similar manner, frequent cleaning, and incubated at 28.5 degrees Celsius. At least 20 embryos for each replicate were collected for both qRT-PCR and RNA-seq. During this study, embryos were periodically cleaned and imaged at 4, 6, 8 hpf and at 1-day post-fertilization (dpf). For knockdown phenotype rescue experiments 20-25 pg of exogenous RNA was used. For α-amanitin experiments, 200 µl/ml α-amanitin was microinjected into one-cell-stage embryos. For 3′UTR reporter assays, 20pg RNA was injected for fluorescence imaging and 10 pg RNA for qRT-PCR analysis.

### Hybridization Chain Reaction (HCR) for zebrafish embryos

Custom probes targeting *cth1*, *tmem243a, lipia*, GFP, and TagRFP transcripts, along with corresponding amplifiers and reagents for V3HCR, were obtained from Molecular Instruments. Hybridization was performed following the manufacturer’s zebrafish embryo protocol. On the second day, amplifiers conjugated to specific fluorophores were added during the amplification step and incubated overnight at room temperature in the dark. Embryos were subsequently washed and counterstained with DAPI prior to imaging.

### RNA isolation and qRT-PCR

To further test the knockdown efficiency and find the molecular signatures of knockdowns, RNA was isolated using either the standard TRIzol method (ThermoFisher Scientific) or the Zymo-Research RNA Mini Kit (ZYMO RESEARCH, Cat. No. R1055).

The concentration of the isolated RNA was measured using either a NanoDrop spectrophotometer or the Qubit RNA-Broad Range assay. For qRT-PCR analysis, complementary DNA (cDNA) was synthesized according to the manufacturer’s protocol using the SuperScript IV First-Strand Synthesis System (ThermoFisher Scientific, Cat. No. 18091050) and using different oligos mentioned in table (Table S1). Quantitative real-time PCR (RT-PCR) was then performed using the low ROX SYBR-Green dye (Quanta Bio, Cat. No. 95074-05K), gene-specific primers (20μM), and 1-2μl of cDNA diluted in a 25μl reaction volume, depending on the gene of interest and the developmental time point. The reactions were conducted on an automated Freedom EVO® PCR workstation (Tecan) using a predefined template program designed for the number of samples. For normalization, endogenous housekeeping genes such as cdk2ap2 or taf15 were used as reference genes for target gene expression analysis, while for reporter assays, other injection controls like TagRFP, and Nano Luciferase were utilized.

### Knockdown phenotype rescue

Guide RNAs (gRNAs) were designed either to target the 3’UTR or the ORF of the *cth1* (Table S1), following a previously published protocol (Hernandez-Huertas et al, 2022). For *cth1* knockdown phenotype rescue experiments, a *cth1* ORF containing eight synonymous mutations at the gRNA target site was cloned with its endogenous 3′UTR to generate a gRNA-resistant transcript (Table S2), enabling transcript expression without interference from ORF-targeting gRNAs. A GFP construct served as a negative control to account for ectopic overexpression effects.

In a complementary rescue strategy, the *cth1* ORF was cloned with a heterologous (non-zebrafish) 3′UTR, thereby avoiding targeting by gRNAs directed against the endogenous *cth1* 3′UTR. As a control, a GFP construct carrying the same heterologous 3′UTR was used to control for both gRNA specificity and overexpression-related effects.

### CRISPR-Cas9 mediated mutation in zebrafish

Three gRNAs were designed using CRISPR Scan guidelines using oligos in (Table S1). The Cas9 plasmid (Addgene # 179317) was digested using *XbaI* enzyme and capped *in vitro* transcription was performed using mMESSAGE mMACHINE™ T3 Transcription Kit to generate Cas9 mRNA (Vejnar et al, 2016). The gRNAs were synthesized according to the protocol outlined for the AmpliScribe-T7 Flash Kit. Zebrafish embryos at the one-cell stage were injected with either 100 pg of Cas9 mRNA and 25 pg of each of the three gRNAs in combination or with 25 pg of each gRNA separately. After injection all the embryos were processed and incubated similarly. Phenotypes of the injected embryos were observed and imaged at 6 hpf and 1 dpf. During this process individual embryos were collected at 24 hpf in QuickExtract™ DNA Extraction Solution (Lucigen) for Indel mutations genotyping using MiSeq2 x 250 flow cell. Sequencing and mutation rates were analyzed as described in genotyping protocol (Billmyre et al, 2023).

### Scanning Electron Microscopy

Sperm samples were diluted onto coverslips coated with 4nm of gold palladium with a Leica EM Ace600 and prepared as previously described for preserving membranes (Korneev et al, 2021). After critical point drying with a Tousimis Samdri-795 and coated with an additional 4nm of gold palladium sperm were imaged at 300kV, 100pA with an SE2 detector on a Zeiss Merlin SEM. All the images were processed similarly in Fiji using CLAHE plugin.

### Gamete collection from Zebrafish

Gametes were collected from wild-type and mutant zebrafish following institutional animal care and handling guidelines. For oocyte collection, females were paired with males overnight in tanks separated by a divider. The following morning, coinciding with light cycle activation, females were anesthetized in 4 g/L MS-222 and stripped to obtain mature oocytes. For sperm collection, males were anesthetized using the same procedure, placed ventral side up in a fish holder, and sperm was collected using a microcapillary pipette connected to an aspiration tube assembly.

### Image acquisition

Different microscopes were used according to the requirement. For zebrafish embryos grouped pictures at early time point up to 1 dpf Leica stereo-fluorescent microscope with DFC900 camera. For individual representative images Leica MZ APO stereo microscope was used for bright field images. For HCR images Nikon spinning disc a confocal microscope was used at 10x and 20x magnification. For large adult fish Samsung galaxy s21 ultra camera was used. For sperm images Zeiss Merlin SEM with SE2 detector was used. For image processing Fiji software was used and image panels were made using adobe illustrator 2024.

### RNA-seq libraries preparation and primary analysis

Total RNA was isolated form 20 embryos using Zymo-Research RNA Mini Kit (ZYMO RESEARCH, Cat. No. R1055) and RNA quality was measured with Bioanalyzer (Agilent). Only RNA with > 7.5 RIN were used for library preparation. For RNA-seq at least 500 ng of RNA was used for library preparation using reagents from Watchmaker Genomics, for total RNA-seq library preparation Watchmaker mRNA Library Prep Kit (Cat. No. 7BK0001) with xGen Stubby Adapter and UDI Primers (IDT, Cat. No. 1000592) were used. Samples were sequenced using paired reads on G4-F3 with a read length of 50bp or 75bp using a flow cell from Singular Genomics (Cat. No. 700125). For Ribosomal RNA depleted RNA-seq, Polaris depletion Kit rRNA/Globin (HMR) (Watchmaker Genomics, Cat. No. 7K0077) was used. Samples were run using AVITI 2×75 Sequencing Kit Cloud break FS High output freestyle flow cell (Element Biosciences, Cat. No. 860-00015). RNA-seq reads were demultiplexed into fastq format allowing up to one mismatch and Singular Genomics sgdemux (v 1.2.0) for samples sequenced on the G4-F3. Subsequently, the reads were aligned to *GRCz11* reference genome from Ensembl using STAR (v 2.7.10b). The gene model retrieved from Ensembl, release 106 was used to generate gene read counts. The transcript abundance TPM (Transcript per Million) was quantified using RSEM (v 1.3.1).

### Western blot

To assess protein expression of *cth1*–α-tagged mRNA used in rescue experiments, zebrafish embryos were injected at the one-cell stage with *cth1*–α-tagged mRNA (10 pg). GFP–α-tagged mRNA (10 pg) was used as a positive control. At 4 hpf, ∼50 embryos per condition (in triplicate) were collected, lysed, and subjected to immunoblot analysis using an anti-α-tag antibody (Nanotag, Cat No: N1581). α-tubulin (Sigma-Aldrich, Cat No: T9026) was used as a loading control.

### RNA immunoprecipitation (RIP) and RT–PCR

To identify Cth1-associated RNAs, zebrafish embryos at the one-cell stage were injected with *cth1*–α-tagged mRNA (20 pg) containing its endogenous 3′UTR. GFP–α-tagged mRNA (20 pg) was used as a control, and tagRFP mRNA (20 pg) was co-injected as an injection control. For each condition, ∼300 embryos per replicate were collected at 4 hpf. Embryos were lysed and subjected to RNA immunoprecipitation using α-tag– specific pulldown (Nanotag: ALFA Selector ST Beads, Cat No. N1516-L), followed by standard washing steps according to the manufacturer’s instructions. A fraction of each lysate (1:100 dilution) was reserved as input. RNA was extracted from both input and pulldown fractions, followed by cDNA synthesis. Enrichment of endogenous Cth1 target transcripts was assessed by RT–PCR. tagRFP amplification was used as a control for injection and specificity.

### Poly(A)-tail status analysis

To assess the polyadenylation status of transcripts, we compared gene expression between poly(A)-enriched and ribo-depleted RNA-seq libraries prepared from the same biological samples. TPM values were obtained from RSEM and averaged across replicates within each sample group. A poly(A) status score was computed for each gene as log2((TPM_polyA + 0.05) / (TPM_ribo + 0.05)), where the pseudo count of 0.05 prevents undefined values for unexpressed genes. Positive scores indicate efficient capture by poly(A) selection (polyadenylated transcripts), while negative scores indicate transcripts predominantly detected by ribo-depletion (non-polyadenylated). To compare poly(A) status changes across experimental conditions, we computed fold-changes in poly(A) status scores between treatment and control samples. Genes with zero TPM in both libraries for a given condition were excluded from downstream comparisons. Cumulative distribution functions (ECDFs) of poly(A) status fold-changes were compared between gene groups of interest and background genes using two-sample Kolmogorov-Smirnov tests, with p-values adjusted for multiple comparisons using the Benjamini-Hochberg method.

### Poly(A)-tail Assay

To assess polyadenylation status of the Cth1 target *tmem243a*, wild-type and mutant *tmem243a* mRNAs (20 pg) were injected into one-cell stage zebrafish embryos. Approximately 50 embryos per condition (in replicates) were collected at 4 hpf. Total RNA was isolated using TRIzol reagent according to the manufacturer’s instructions. An RNA adaptor was ligated to the 3′ end of the purified RNA, followed by cDNA synthesis using an oligo complementary to the adaptor sequence. cDNA was subsequently purified using a Zymo Clean & Concentrator kit. PCR amplification was performed to specifically detect injected *tmem243a* transcripts poly(A)-tail status using a forward primer targeting the injected RNA and a reverse primer complementary to the adaptor-derived sequence.

### Construction and GFP:3′UTR reporter assay

To generate a GFP:3′UTR individual reporter, 3′UTRs of candidate targets identified from RNA-seq were amplified from cDNA prepared from 1 hpf zebrafish embryos using oligos in (Table S1). Amplified fragments were cloned downstream of GFP and sequence verified. Positive clones were digested with KpnI-HF, and capped RNAs were synthesized *in vitro* using the SP6 mMESSAGE mMACHINE Maxiscript kit. For qRT-PCR assays, an RNA cocktail was prepared containing six different GFP:3′UTR reporter transcripts (10 ng/µl each), GFP: *β-globin* 3′UTR, TagRFP (10 ng/µl), NanoLuc (10 ng/µl), and RfxCas13d mRNA (300 ng/µl). For *cth1* knockdown, gRNA (600 ng/µl) targeting *cth1* was co-injected with the cocktail. Embryos were collected at 6 hpf for RNA extraction, cDNA synthesis, and qRT-PCR. For fluorescence imaging, each GFP:3′UTR reporter was injected individually (20 ng/µl) with TagRFP control, and embryos were imaged at 8 hpf.

### Construction of *cth1* GFP:3′UTR reporter library

Cth1 3′UTR fragments of 50 nucleotides each with 2-5 nucleotides overlap with consecutive fragments along with fragments from miR-430 target sod1 genes as a positive control downstream to GFP reporter were ordered from IDT. The whole reporter and 3′UTR fragments cassette was flanked with illumina adaptors. Whole library was minimally PCR amplified (20 cycles) using a set of universal oligos, ran on agarose gel and desired size band was cut, gel extraction purification was done, and in vitro transcription was performed using the SP6 mMESSAGE mMACHINE Maxiscript kit. The library (15 pg/embryo) was injected at one cell stage embryo and 50 embryos each were collected in triplicate at 2 and 8 hpf for RNA isolation. RNA was poly(A) enriched using Dynabeads™ mRNA Purification Kit (Thermofisher, Cat No. 61006) and sequenced.

### Data Analysis

Data analysis was carried out in R (version = 4.4.1) with packages from CRAN and Bioconductor. RNA-seq data were imported and filtered using tidyverse and data.table, retaining protein-coding genes with moderate expression (CPM >= 5 at 2hpf) and 3′UTRs between 100 and 800 nt. Differential expression results from RfxCas13d knockdowns were merged, and a putative cth1 target group was defined by requiring log2fold-changes ≥ 0.5 and adjusted P < 0.05 under both g1CTH1 and g2CTH1 conditions, while four control sets were generated by randomly sampling non-responsive genes (C1-C4). Gene-group assignments were visualized with scatter plots created using ggplot2 and ggpubr. Time-course expression data from ribosomal depletion, poly(A) selection and α-amanitin-treated libraries were log2-transformed and plotted (Medina-Munoz et al, 2021). Median expression was overlaid with cross-bars, and RNA-decay rates were estimated as log2 ratios of late to early timepoints. Differences between candidate and control groups were assessed with Wilcoxon rank-sum tests and visualized via boxplots and distribution curves. Transcript annotations, including coding sequence and UTR lengths, were retrieved from Ensembl via biomaRt. Sequences were manipulated with Biostrings. K-mer counts in candidate versus control 3′UTRs were calculated with an in-house script and compared using one-sided Fisher’s exact tests, with Benjamini– Hochberg correction; per-k-mer P-values were combined across control comparisons using Fisher’s combined probability test. Enriched motifs were illustrated with sequence logos, and their transcript positions were mapped back onto 3′UTRs. Distances from motifs to poly(A)-tails were calculated and plotted to infer positional bias. The source code for the analysis presented in this study is available from GitHub.

### Quantification and Statistical analysis

To select different sample sizes, no predefined statistical method was used. The experiments were not randomized, and investigators were not blinded to group allocation during the experiments or outcome assessment. All collected data were included in the analysis without exclusion. All the quantification, gene expression and RNA-Seq plots were generated in R. Phenotype quantification across different CRISPR-Cas-mediated injection conditions was analyzed using chi-squared (χ²) tests, performed using R. For quantitative RT-PCR analysis, knockdown rescue experiments, p-values were determined using an unpaired t-test, and error bars represent the Standard Error of the Mean (SEM) with statistical analysis performed on at least two biological replicates. P value for cumulative plots between different sets of genes was calculated using Wilcoxon rank-sum tests.

## Lead Contact

Further information and resource request should be directed to lead contact Ariel A. Bazzini.

## Data and Code Availability

The Accession number for the RNA-Seq data used in this manuscript is GEO: GSE330436. All the relevant data is available upon request from the corresponding author.

### Declaration of generative AI and AI-assisted technologies in the writing process

During the preparation of this work the authors used ChatGPT in order to improve only the text. After using this tool, the authors reviewed and edited the content as needed and take full responsibility for the content of the publication.

## Acknowledgments

We thank Robb Krumlauf, Anthony Treichel, Ismael Moreno Sanchez, Miguel Moreno-Mateos for critical reading and valuable suggestions for this manuscript. Carrie Carmichael for zebrafish related discussion and training. Different core facilities at Stowers Institute like Sequencing and Discovery Genomics, genome engineering, automation and PCR technology, Media prep and all members of the Bazzini laboratory for their intellectual and technical support. This study was supported by the Stowers Institute for Medical Research. AAB was awarded a Pew Innovation Fund and with the US National Institutes of Health (NIH-R01 GM136849, NIH-R21 OD034161 and NIH-R35 GM161459). This work was performed as part of post-doctoral training for GK at the Stowers Institute for Medical Research. This work was performed as part of thesis research for D.B.A. at the Graduate School of the Stowers Institute for Medical Research. Original data underlying this manuscript can be accessed from the Stowers Original Data Repository at http://www.stowers.org/research/publications/libpb-2587.

## Author contributions

G.K. and A.A.B. conceived the project and designed the research.

1. G. K. performed all the experiments.

H.H. did primary RNA-seq analysis.

G.K, DAB, DBA and A.A.B. performed data analysis.

S.N. performed electron microscopy.

G.K, and A.A.B. wrote the manuscript with input from the other authors.

All authors reviewed and approved the manuscript.

---

**Supplementary Figure 1.**
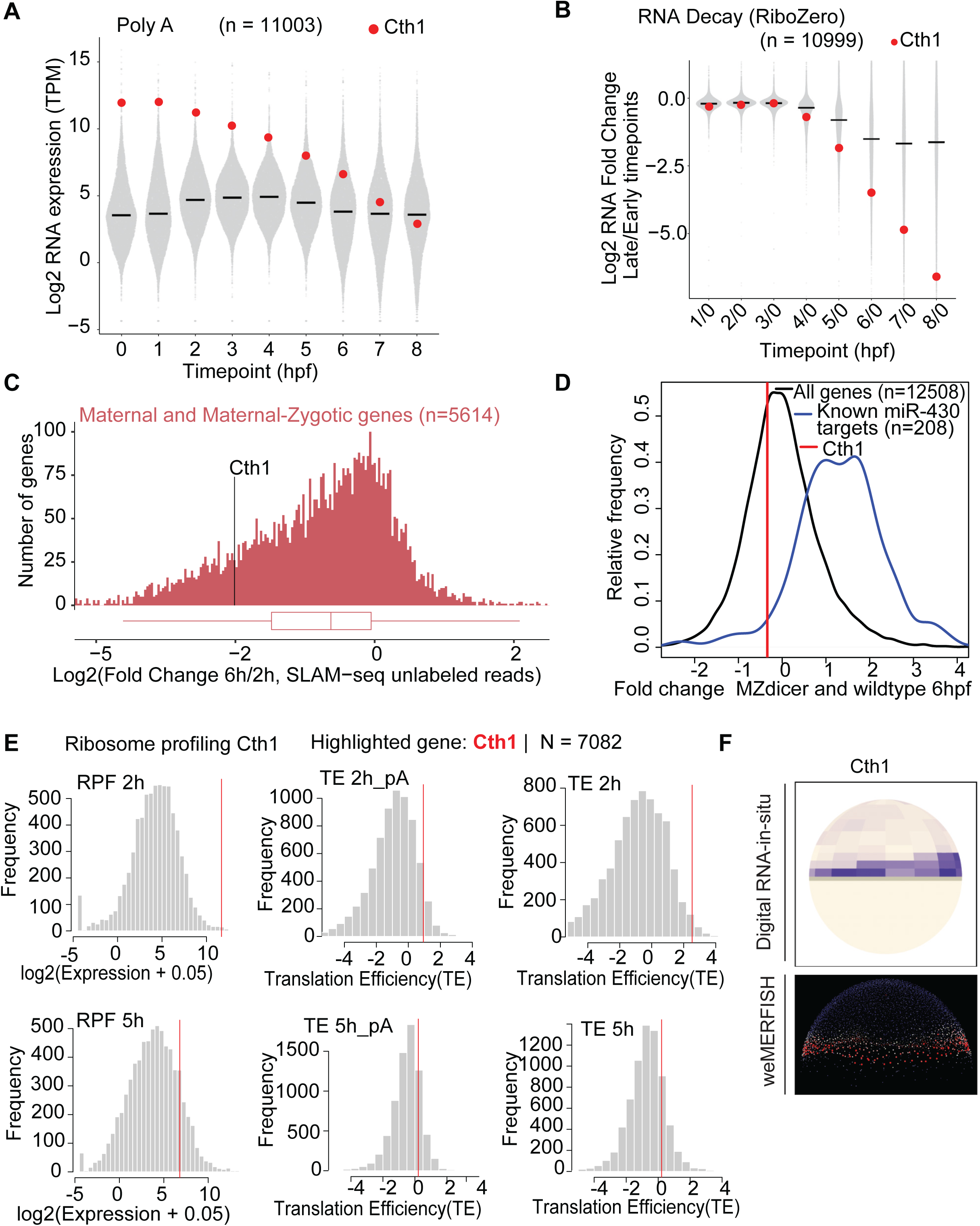
*Cth1* is one of the most highly maternally deposited mRNAs but is rapidly degraded during the maternal-to-zygotic transition. A. PolyA-enriched RNA-seq time course (0-8 hpf) showing cth1 mRNA is among the top maternally deposited (top 75) mRNA at 0 hpf while getting rapidly degraded with time by 8 hpf. Data is derived from SG Medina-Muñoz et al., *Genome Biology,* 2021. B. mRNA decay measured as fold change between Ribosomal RNA–depleted RNA-seq at 0 hpf and indicated time point (0-8 hpf) showing that cth1 mRNA is one of the most unstable mRNA (top 20). Data is derived from SG Medina-Muñoz et al., *Genome Biology,* 2021. C. SLAM-seq data analysis measuring the log fold change of maternal and maternal zygotic transcripts level between 6 hpf and 2 hpf. SLAM-seq data further validate the pronounced instability of cth1 among 5,614 purely maternal and maternal–zygotic transcripts. Data is derived from DB Amaral et al., *Genome Biology,* 2024. D. Histogram showing the relative abundance of *cth1* transcripts at 6 hpf in MZ dicer mutants compared to wild-type embryos, indicating that *cth1* regulation is independent of miR-430 activity. In black line are all the genes and in blue genes known to be miR-430 targets. n indicate the number of genes in each group. E. Ribosome profiling analysis of *cth1* translational activity at 2 hpf and 5 hpf. Histograms show genome-wide distributions of ribosome-protected fragment levels, poly(A)-normalized translation efficiency, and total RNA-normalized translation efficiency, with cth1 indicated by a red line. cth1 shows high ribosome occupancy and elevated translation efficiency at both stages. Data is derived from Bazzini et al., *EMBO Journal*, 2014. F. Digital RNA *in-situ* and weMERFISH showing the predicted and calculated *cth1* RNA marginal distribution respectively at mid blastula stage in zebrafish embryo (Satija, Farrell et al., Nature Biotechnology, 2015, Yinan Wan et al., Science, 2026).

**Supplementary Figure 2.**
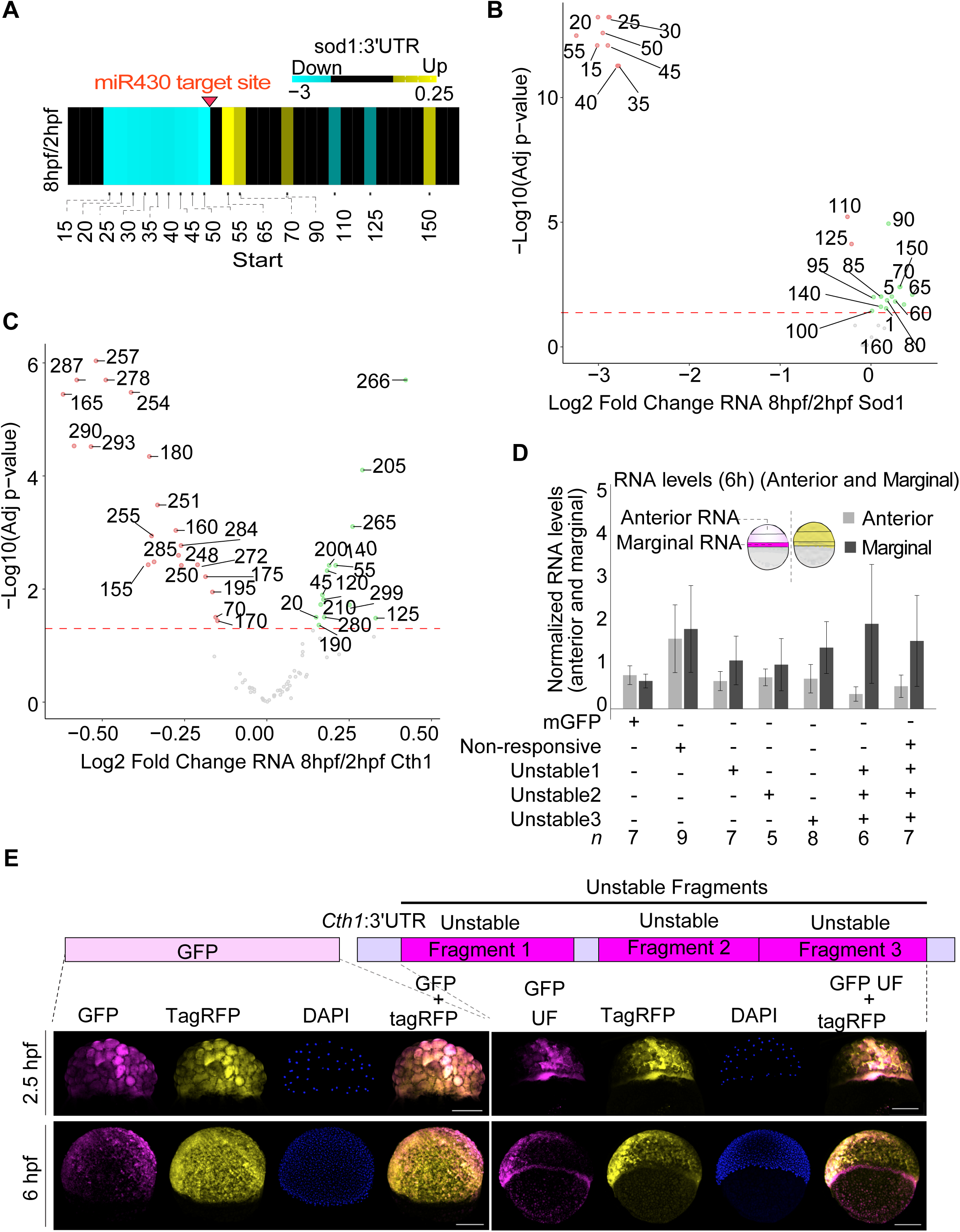
The *cth1* 3′UTR harbors *cis*-regulatory elements driving marginal localization. A. Heat map showing the stability of different *sod1* 3’UTR fragments (known miR430 target) between 8 and 2 hpi. Cyan color indicates unstable regions, black color non-responding region and yellow stable regions. The number represent the 5’position of the fragment with respect to *sod1* 3’UTR. Red color line indicates the fragments that contain the full miR430 binding site. B. Volcano plot showing the *sod1* 3′UTR fragments levels as the log fold change between 8 and 2 hpf. Unstable fragments are highlighted in red, while non-responding fragments are shown in green dots. The numbers represent the 5’position of the fragment in the *sod1* 3′UTR. C. Volcano plot showing the *cth1* 3′UTR fragments levels as the log fold change between 8 and 2hpf. Unstable fragments are highlighted in red, while non-responding fragments are shown in green dots. The numbers represent the 5’position of the fragment in the *cth1* 3′UTR. D. Bar plot showing normalized HCR RNA quantification for GFP normalized with TagRFP RNAs for two regions of the same embryos (anterior and marginal) for different *cth1* 3′UTR fragments and controls. Reporters carrying unstable cth1 3′UTR fragments show reduced anterior RNA levels, and enriched marginal region patterning while controls RNAs remain uniformly distributed. Light grey and dark grey bars are for anterior and marginal regions respectively. *N* is number of embryos analyzed. E. GFP and TagRFP HCR showing that all injected mRNAs were uniformly distributed at 2.5 hpi, but at 6 hpi only GFP-containing the full unstable *cth1*-3’UTR fragment (unstable fragments 1+2+3, 175 ntd) identified by the tilling library enriched to the marginal region, while the GFP and other injected control TagRFP mRNA display uniform distribution across the embryo. Scale bar, 200 μm.

**Supplementary Figure 3.**
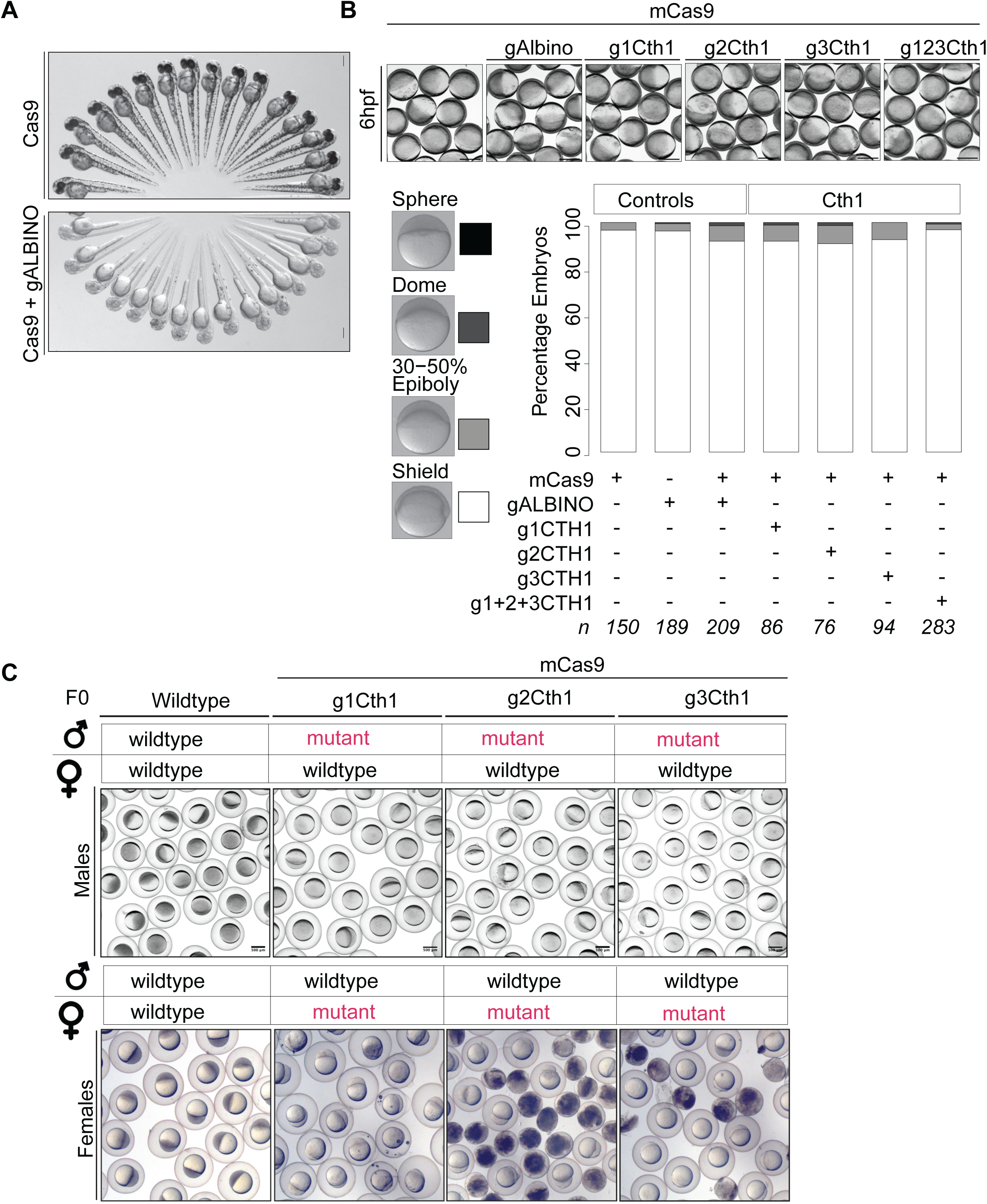
*Cth1* F0 mutants do not exhibit early developmental defects but display severe infertility. A. Images of 2 dpf zebrafish larvae (SpyCas9 and SpyCas9+gALBINO) showing loss of pigmentation in *albino* mutant as compared to control. Scale bar, 2mm. B. Representative images and stack bar plot showing that no major developmental phenotypes were observed from F0 embryos injected with any of the guideRNAs and/or Cas9 combination at 6hpi. Different intensities of gray color are directly associated with severity of developmental phenotype (sphere (black) to shield (white)). Scale bar, 0.5mm. C. Representative embryos images after crossing wildtype and/or SpyCas9 *cth1* F0 mutants (red color text) adults from three independent gRNAs. Mutant females produced either dead or unfertilized embryos, mutant males produced unfertilized embryos, while wildtype crosses produced viable embryos. (Panels show 4 hpf embryos). *n* represents the number of embryos observed, Scale bar, 1000 μm.

**Supplementary Figure 4.**
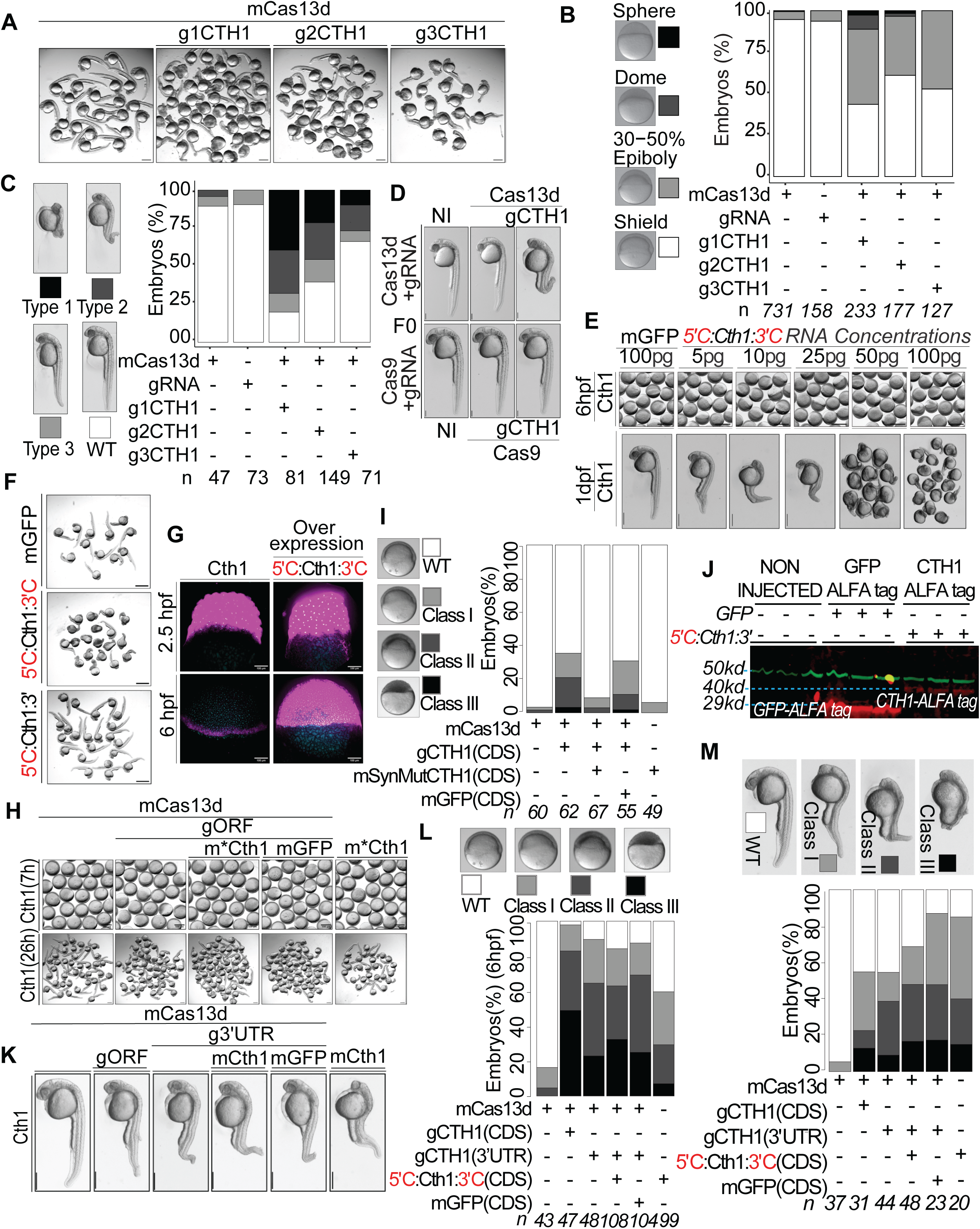
Cas13d-mediated knockdown reveals the essential role of *cth1* during the maternal-to-zygotic transition. A. RfxCas13d mediated *cth1* mRNA knockdown phenotypes images at 1 dpf along with RfxCas13d alone controls using three independent gRNAs (g1CTH1, g2CTH1 and g3CTH1). Scale bar, 1 mm. B. Stacked bar plot showing the percentage of developmentally affected embryos for *cth1* mRNAs knockdown along with controls (RfxCas13d and gRNA alone) at 6 hpf. The intensity of gray color is directly associated with severity of the developmental phenotypes (sphere, dome, 30-50% epiboly, shield) respectively. n represents the number of embryos analyzed. C. Stacked bar plot is showing the percentage of developmentally affected embryos from three independent gRNAs for *cth1* mRNAs along with controls (RfxCas13d and gRNA alone) at 1 dpf. The intensity of gray color is directly associated with severity of the developmental phenotypes (Type 1(black), to wildtype (white)) respectively. n represents the number of embryos analyzed. D. Comparison of representative 1 dpf zebrafish embryo images of RfxCas13d mediated *cth1* mRNA knockdown and SpyCas9 mediated F0 *cth1* gene mutants. Scale bar, 1mm. E. Images at 6 hpf and 1 dpf showing effect of mRNA encoding for *cth1* with control 5’ and 3’UTRs. Overexpression induces body axis defects even at the lowest concentration tested (5 ng/µl) as compared to control GFP with control 5’ and 3’UTRs. Scale bar, 1 mm. F. Images at 1 dpf showing that injection of mRNA encoding for *cth1* with control 5’ and 3’UTRs (25 ng/ul) affected development while injection of *cth1* mRNA with its endogenous 3’UTR (25ng/ul) does not affect development compared to embryos injected with mRNA encoding for GFP. Scale bar 1mm. G. HCR for *cth1* mRNA in un-injected and injected embryos with *cth1* ORF alone (10 ng/µl) at 2.5 hpf and 6 hpf. At 2.5 hpf, *cth1* expression is uniform across both conditions. By 6 hpf, un-injected embryos display characteristic marginal enrichment of *cth1* mRNA, whereas embryos injected with *cth1* ORF alone retain uniform distribution, indicating that the *cth1* 3′UTR is required for proper transcript localization. Magenta, *cth1* mRNA; cyan, nuclei (DAPI). Scale bar, 100 µm. H. Images at 6 hpf and 1 dpf showing the developmental phenotype associated with *cth1* knockdown (RfxCas13d + gORF), interestingly, the phenotype was rescued by ectopic expression of *cth1* insensitive to the guideRNA (RfxCas13d + gORF + *m*cth1(synMut)*) but not by the ectopic expression of GFP (RfxCas13d + gORF + mGFP). Injection of RfxCas13d alone or RfxCas13d + gORF and m*\*cth1(synMut)* mRNAs did not affect development. Images are of 6 hpf and 28 hpf embryos, scale bar, 500 µm. I. Stack bar plots showing the percentage of observed phenotype rescue for *cth1* (at 6 hpf. Different intensities of gray color are positively associated with the severity of the phenotype (dark, most severe phenotype (class III) and white, (wildtype)). *n* represents the number of embryos observed in each condition. J. Western blot analysis of ALFA-tagged proteins in zebrafish embryos at 4 hpf. Embryos were injected with mRNAs encoding GFP–ALFA tag (positive control) or Cth1–ALFA tag, and non-injected embryos as a negative control. ALFA-tagged proteins (Red) were detected at the expected sizes for GFP–ALFA (∼40 kDa) and Cth1–ALFA (∼29 kDa); α-tubulin (Green) served as a loading control. K. Representative brightfield images of a Cas13d-mediated *cth1* knockdown and rescue assay in zebrafish at 26 hpf. For the rescue experiment, the endogenous *cth1* 3′UTR was targeted by a specific guide RNA (gCTH1-3′UTR), while an ectopic *cth1* mRNA carrying a non-targeted 3′UTR (mCTH1-CDS) was co-injected to restore Cth1 protein levels. Additional controls include: Cas13d alone, Cas13d with a guide RNA targeting the *cth1* coding sequence (gCTH1-CDS), Cas13d with gCTH1-3′UTR plus mGFP mRNA, and *cth1* ORF mRNA alone. Notably, ectopic *cth1* mRNA carrying a non-targeted 3′UTR failed to rescue the knockdown phenotype, indicating that the *cth1* 3′UTR is required for function. L. Stacked bar graphs showing the percentage of developmentally affected embryos at 6 hpf following injection of ectopic *cth1* ORF mRNA (10 ng/µl) or GFP mRNA as a control (10 ng/µl). Representative brightfield images illustrate the range of developmental phenotypes classified as wild type (Class I, white), mildly affected (Class II, grey), or severely affected (Class III, black). Darker shading reflects increasing severity of developmental defects. *n* indicates the number of embryos scored per condition. M. Stacked bar graphs showing the percentage of developmentally affected embryos at 26 hpf following the same injection conditions as in C. Representative brightfield images show developmental phenotypes at 26 hpf classified by the same scheme. *n* indicates the number of embryos scored per condition.

**Supplementary Figure 5.**
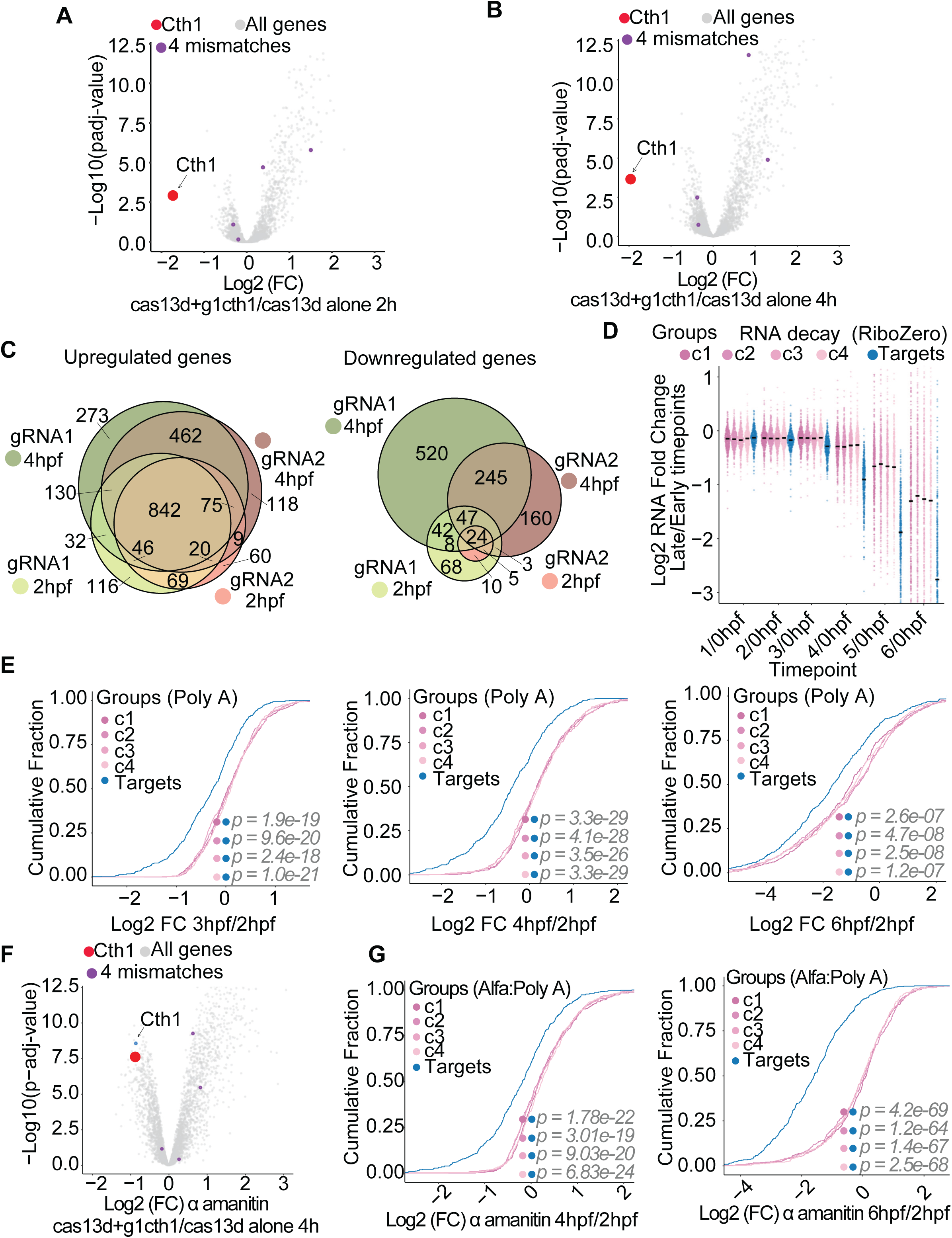
*Cth1* regulates mRNA stability during the MZT via 3′UTR *cis* elements. A.B. Volcano plots showing the mRNA fold change in embryos co-injected with RfxCas13d and g1CTH1 guideRNAs compared to embryos injected with RfxCas13d alone at 2 (A) and 4 (B) hpf using g1CTH1. *Cth1* was efficiently knockdown (red) and no off targets were identified allowing up to 4 mismatches (purple dots) between g1CTH1 and downregulated gene sequences which suggest specific *cth1* mRNA knockdowns. C. Ven diagram showing overlap of upregulated and downregulated genes following *cth1* knockdown with g1CTH1 and g2CTH1 at 2 and 4 hpf. A significant number of upregulated genes show overlap with both gRNAs’ knockdowns, supporting high knockdown specificity (2 hpf g1CTH1 vs. g2 CTH1: p value ≤ 0.2e-16; 4 hpf g1 CTH1 vs. g2 CTH1: p value ≤ 0.2e-16; Fisher’s Exact Test). D. Sina plot showing the log fold change of transcripts levels for ribosomal RNA–depleted RNA-seq during embryogenesis (1 to 6 hpf) compared to the 1 cell stage embryo (0 hpf). Potential *cth1* targets are shown in blue and the control groups (C1-C4) in shades of different pink color. E. Cumulative fraction PolyA RNA-seq plot showing fold change between indicated hpf (3, 4 ,6 hpf) and 2 hpf. The putative Cth1 targets (blue) are unstable as compared to control groups (C1-C4, pink color). Wilcoxon rank-sum test, P value ≤ 1.2e-07. F. Volcano plots showing the mRNA fold change in α-amanitin treated embryos co-injected with RfxCas13d and g1CTH1 compared to embryos injected with RfxCas13d alone at 4 hpf using g1CTH1. *Cth1* was efficiently knockdown (red) and no off targets were identified allowing up to 4 mismatches (purple dots) between g1CTH1 sequence and downregulated gene sequences which suggest specific *cth1* mRNA knockdowns even in α-amanitin treated embryos. G. Cumulative fraction in PolyA RNA-seq plot from α-amanitin treated embryos showing fold change between 4 and 2 hpf, 6 and 2 hpf. The putative Cth1 targets (Blue) are unstable as compared to control groups (C1-C4, pink color). Wilcoxon rank-sum test, P value ≤ 3.01e-19.

**Supplementary Figure 6.**
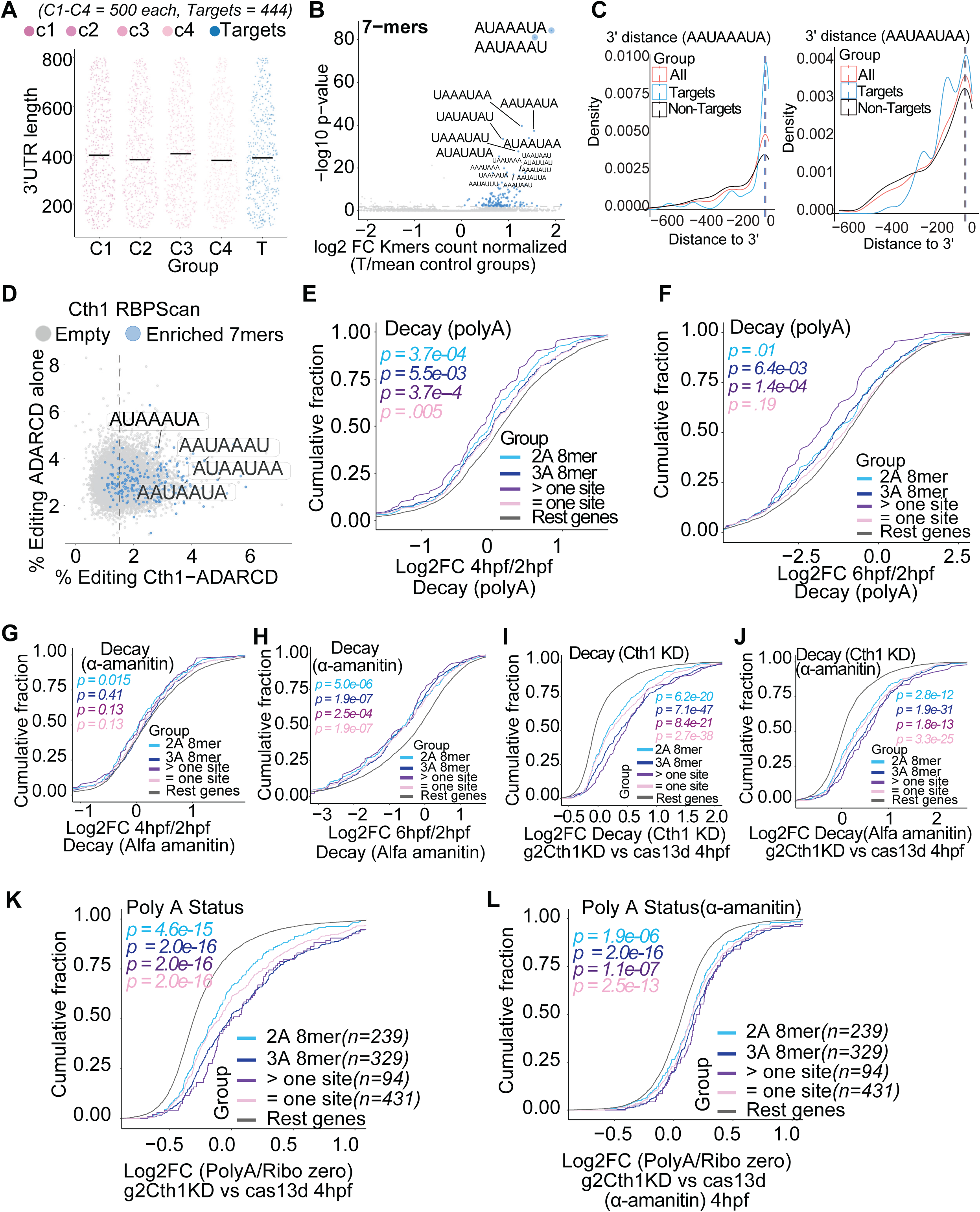
Cth1 acts on the AU -rich 3′UTR of its targets. A. Sina plot showing that there is no difference in the 3’UTR length of randomly selected 4 groups (C1-C4, pink color, 500 genes in each group) and the putative Cth1 targets (444 genes). P value ≥ 0.5. B. Scatter plot showing 7-mer motifs enrichment/depletion between the putative Cth1 target group and the 4 controls groups (C1-C4). AU-rich motifs are particularly overrepresented, with sequences containing “AAA” and “AA” showing the strongest enrichment when compared using log fold-change of normalized k-mers. C. The 8-mer motifs containing “AAA” (e.g., AAUAAAUA) show positional bias for the putative Cth1 target group, being enriched within the last 30 nucleotides of 3’UTRs. In contrast, simple “AA” motifs do not exhibit this bias. D. Scatter plot displaying RBPscan data (Kretov, Sanborn et al. 2025), an *in vivo* RNA binding method to identify sequences recognized by the over-expression of an RBP fused to ADAP. The 7-mers motifs (AU rich motifs) enriched with the potential Cth1 target group were also showing Cth1 binding activity by RBPscan. E. Cumulative fraction PolyA RNA-seq plot showing fold change between 4 and 2 hpf (E), 6 and 2 hpf (F). Transcripts containing enriched 7-mers/8-mers, particularly those with “AAA” and multiple motifs, exhibit greater destabilization than other genes. Wilcoxon rank-sum test, P value ≤ 0.005(E), P value ≤ 0.19(F). G. Cumulative fraction in PolyA RNA-seq plot from α-amanitin treated embryos showing fold change between 4 and 2 hpf (G), 6 and 2 hpf (H). Transcripts containing enriched 7-mers/8-mers, especially those with “AAA” and multiple motifs, exhibit greater instability than other genes. Wilcoxon rank-sum test, P value ≤ 0.41(G), P value ≤ 2.2e-04 (H). I.J. Cumulative fraction in PolyA RNA-seq plot upon *cth1* knockdown mediated by g1CTH1 showing fold change between 4 and 2 hpf (I) and α-amanitin treated embryos (J). Transcripts containing enriched 7-mers/8-mers, especially those with “AAA” and Multiple motifs, exhibit more stability than other genes. Wilcoxon test, P value ≤ 6.2e-20 (I), Wilcoxon rank-sum test, P value ≤ 2.8e-12 (J). K. Cumulative distribution plot of poly(A) status, as the log2 fold change between poly(A)-selected and ribo-depleted RNA-seq, upon *cth1* knockdown by g2Cth1KD versus Cas13d control at 4 hpf. Transcripts containing enriched 7-mers/8-mers, especially those with “AAA” and multiple motifs, show significantly greater poly(A) tail enrichment relative to other genes. Wilcoxon rank sum test, number of genes (*n*) and *p*-values are indicated. L. As in K, but in α-amanitin–treated embryos using guide RNA g2 (g2Cth1KD), further validating the finding that *cth1* regulates poly(A) tail length of maternal transcripts containing adenosine-rich 7-mers/8-mers motifs.

**Supplementary Figure 7.**
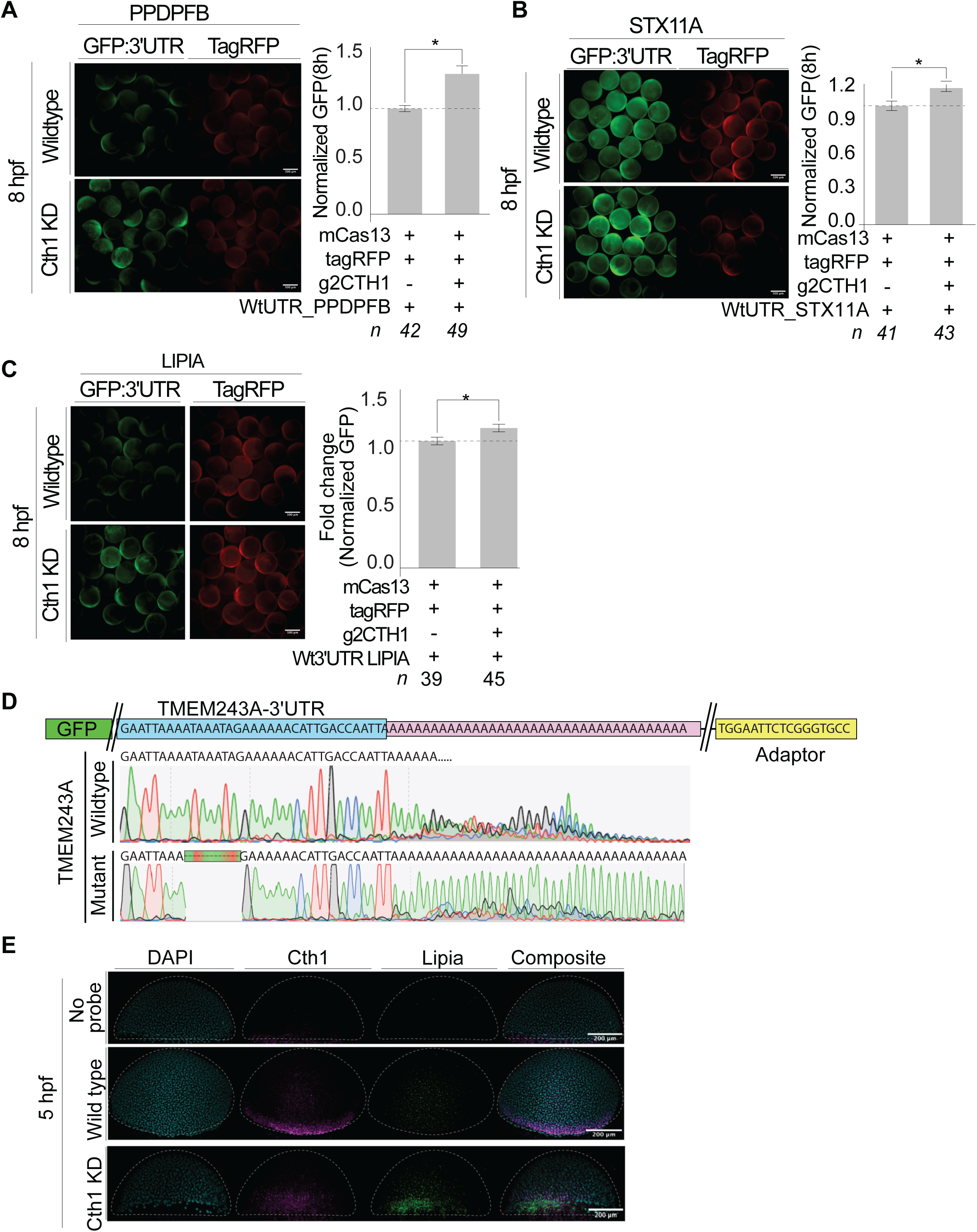
Cth1 acts on the AU - rich 3′UTR of its targets and restricts their spatial patterns. A. Embryos showing stable GFP fluorescence in GFP: 3′UTR*ppdpfb* reporter– injected embryos upon *cth1* knockdown as compared to wildtype conditions at 8 hpf. TagRFP was co-injected as a control. Bar plot showing the normalized GFP quantification in wildtype and *cth1* knockdown conditions. “n” number of embryo calculated, and significance was determined by *t*-test (*p* < 0.05). B. Embryos showing stable GFP fluorescence in GFP:3′UTR*stx11a* reporter–injected embryos upon *cth1* knockdown as compared to wildtype conditions at 8 hpf. TagRFP was co-injected as a control. Bar plot showing the normalized GFP quantification in wildtype and *cth1* knockdown conditions. “n” number of embryo calculated, and significance was determined by *t*-test (*p* < 0.05). C. Embryos showing stable GFP fluorescence in GFP:3′UTR*lipia* reporter–injected embryos upon *cth1* knockdown as compared to wildtype conditions at 8 hpf. TagRFP was co-injected as a control. Bar plot showing the normalized GFP quantification in wildtype and *cth1* knockdown conditions. “n” number of embryo calculated, and significance was determined by *t*-test (*p* < 0.05). D. Sanger sequencing of the RT-PCR bands isolated in (Figure 7H), confirming the identity of the amplified products. The sequence chromatograms reveal a longer poly(A) tail in the mutant reporter compared to the wild-type reporter, consistent with reduced deadenylation in the absence of Cth1 binding motifs. E. HCR analysis of endogenous *cth1* and *lipia* mRNA expression in wild-type and *cth1* knock-down (Cth1 KD) zebrafish embryos at 5 hpf. In wild-type embryos, *cth1* mRNA (Magenta) is enriched at the embryonic margin, while *lipia* mRNA (Green) is lowly expressed. Upon *cth1* knockdown, *lipia* mRNA (Green) exhibits markedly enhanced marginal localization, suggesting that Cth1 restricts the marginal enrichment of *lipia* transcripts. No-probe embryos were included as negative controls to confirm signal specificity. Cyan: DAPI (Nuclei), rightmost panel: Composite images, Scale bar, 200 μm.

